# METAHICT enables comprehensive genome-resolved microbiome analysis with metagenomic Hi-C

**DOI:** 10.1101/2025.10.12.681839

**Authors:** Shiyuan Wang, Zhen Qin, Hang Yu, Ruishan Liu, Yong Ge, Maitreya Dutta, Luan Vu, Yuxuan Du

## Abstract

Metagenomic Hi-C (metaHi-C) adds within-cell proximity information to shotgun sequencing, but existing analyses are fragmented and short-insert intra-contig read pairs from Hi-C libraries are often underused. Here, we present METAHICT, a modular and configurable workflow spanning read processing, metagenome-assembled genome (MAG) reconstruction, and downstream analysis. METAHICT consolidates outputs from bin3C, MetaCC, and ImputeCC and uses an expectation–maximization model of mapped insert sizes to select short-insert, shotgun-like Hi-C pairs for per-bin reassembly. Across seven datasets, binning and consolidation generally recovered more near-complete and medium-quality MAGs than individual Hi-C binners. On the sheep-gut long-read assembly, METAHICT’s binning and consolidation stage recovered 929 MAGs meeting at least the medium-quality threshold, including 487 near-complete MAGs. Within the reassembly module, adding EM-selected Hi-C reads to shotgun-only inputs yielded further significant reductions in contamination across multiple environments. The workflow also integrates scaffolding, taxonomic annotation, contact-map visualization, and analysis of candidate mobile genetic element (MGE)–host pairs. Together, these capabilities provide standardized outputs for comparative and hypothesis-driven genome-resolved microbiome analysis.

## 1 Introduction

Metagenomics profiles microbial communities directly from environmental or host-associated samples without isolating or culturing individual taxa, enabling analyses of community structure and metabolic potential in situ [1–4]. The recent integration of high-throughput chromosome conformation capture (Hi-C) with whole-metagenome shotgun (WMG) sequencing has added a spatial dimension to these studies: by recording physical co-localization of DNA within intact cells, metagenomic Hi-C (metaHi-C) provides long-range linkage that can associate fragments originating from the same cell and inform downstream genome-resolved analysis [5–10]. In a typical metaHi-C workflow, shotgun sequencing extracts and sequences DNA fragments from a single microbial sample, while a parallel Hi-C protocol on the same material crosslinks chromatin, performs proximity ligation to join spatially co-located loci, and yields paired-end reads capturing within-cell contacts. In this context, metaHi-C datasets can be categorized by the sequencing strategy of the shotgun library: short-read metaHi-C (e.g., Illumina) versus long-read metaHi-C (e.g., PacBio or Nanopore). Shotgun reads are assembled de novo into contiguous sequences, termed contigs, and Hi-C read pairs are then aligned to these contigs. The number of Hi-C pairs that bridge any two contigs defines a contig-by-contig contact matrix that reflects their relative spatial proximity within the cell. Because raw contact counts are influenced by confounding factors unrelated to within-cell proximity, such as contig length and coverage, restriction-site density, GC content, and mappability, normalization is required before quantitative interpretation [11–15]. Once appropriately normalized, metaHi-C contacts supply long-range signal, which can be used to group contigs into metagenome-assembled genomes (MAGs) [16] via Hi-C–guided binning and to link mobile genetic elements (e.g., viruses and plasmids) with their microbial hosts. These capabilities have enabled the construction of large genome compendia from complex communities and opened avenues to study species diversity, ecological interactions, and MGE–host relationships within single samples.

Routine metaHi-C analysis remains technically demanding. A typical workflow spans adapter and chimeric-read handling, read mapping, contact construction, bias mitigation, binning, and downstream interpretation, each step invoking different tools with distinct dependencies and file formats [17–20]. In practice, investigators must reconcile library/version conflicts, convert intermediate outputs with custom scripts, and tune parameters that are reported inconsistently across studies. This fragmentation raises the barrier to entry and complicates reproducibility, especially for laboratories without dedicated computational support. Genome-resolved analysis with metaHi-C seeks to recover metagenome-assembled genomes (MAGs) informed by Hi-C contacts and to evaluate their quality by completeness and contamination. Multiple Hi-C–aware binners exist, but none performs consistently best across habitats and assemblies, making principled refinement of results from multiple methods a practical necessity [21]. At the same time, most pipelines prioritize inter-contig contacts and underuse informative intra-contig pairs that could be systematically recruited to strengthen the MAG reconstruction process. Beyond genome recovery, support for downstream steps is limited: MAG scaffolding from contact maps, standardized taxonomic and abundance summaries, Hi-C-based MGE-host linking, and integrated visualization are not well served by current tools. Therefore, an integrated and rigorously benchmarked workflow is needed to combine complementary binning results, make better use of sequence information in Hi-C libraries, and connect genome recovery with downstream analyses.

Here, we introduce METAHICT (METAgenomic HI-C Toolbox), a modular and configurable workflow for end-to-end metaHi-C analysis. Implemented in Nextflow DSL2 [22] with versioned software environments, METAHICT connects raw-read preprocessing, assembly and alignment, Hi-C contact construction and normalization, genome retrieval, and downstream analyses while managing software dependencies and recording analytical settings. For MAG reconstruction, the workflow consolidates outputs from multiple Hi-C-aware binners and fits a mixture model to mapped insert sizes using expectation–maximization (EM) [23] to select short-insert, shotgun-like intra-contig read pairs from Hi-C libraries for per-bin reassembly. The workflow also supports Hi-C-guided scaffolding, taxonomic and abundance profiling, contact-map visualization, and MGE–host analysis.

We evaluate METAHICT on seven datasets spanning host-associated microbiomes, including human gut [13], sheep gut [24], pig gut [25], cow rumen [26], and bovine skin [27], as well as the environmental wastewater [10] and hydrothermal-mat [28] microbiomes. The binning benchmark includes all seven datasets, including the two long-read assemblies, whereas EM-based read selection and per-bin reassembly are evaluated on the five short-read datasets for which the reassembly module is designed. Across datasets, METAHICT generally increases the recovery of near-complete and medium-quality MAGs relative to established Hi-C–based baselines. The per-bin reassembly procedure reduced contamination, and paired ablation showed that adding the EM-selected short-insert read pairs produced further significant contamination reductions relative to shotgun-only reassembly. Bray–Curtis dissimilarities [29] of METAHICT-derived community profiles recapitulated habitat-specific structure, while the recovered MAGs reveal biologically informative signals, including several *Erysipelotrichales* MAGs without a species-level assignment by GTDB-Tk in the sheep gut. A focused case study shows that Hi-C–guided scaffolding improves contiguity for an abundant *Bacteroides vulgatus* MAG, and MGE analysis reports filtered candidate virus–host pairs involving *Faecalibacterium* MAGs. Taken together, these results demonstrate that METAHICT standardizes metaHi-C analysis across protocols and sequencing modalities and provides integrated outputs for comparative and hypothesis-driven genome-resolved microbiome studies.

## 2 Results

### 2.1 Overview of METAHICT

METAHICT (i) processes raw metagenomic shotgun and Hi-C reads to generate assemblies, alignments, coverage profiles, raw and normalized Hi-C contact matrices, and quality-control summaries; (ii) integrates three Hi-C-based binners, including bin3C [17], MetaCC [20], and ImputeCC [30], and consolidates their bin sets into a single, non-redundant MAG collection; (iii) models the mapped insert-size distribution using a mixture model fitted by expectation–maximization (EM), selects short-insert, shotgun-like intra-contig read pairs from Hi-C libraries, and includes these reads alongside matched shotgun reads in per-bin reassembly; and (iv) provides standardized interfaces to established downstream tools, including YaHS for Hi-C-guided scaffolding [31], GTDB-Tk for taxonomic assignment [32], and geNomad for MGE identification [33] (Fig. 1).

**Fig. 1:**
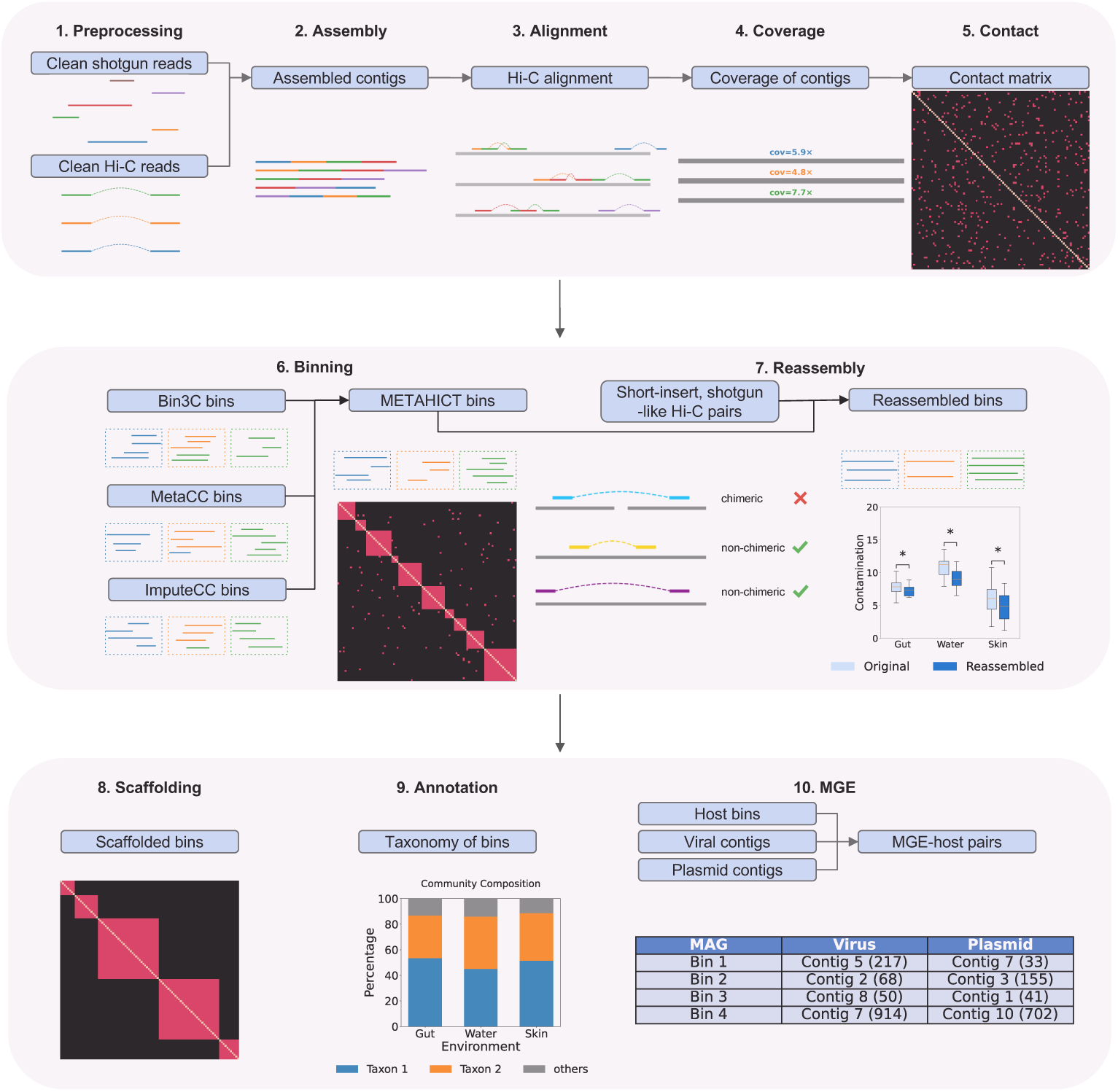
Overview of METAHICT. Inputs are a shotgun library, either short- or long-read, and a matched Hi-C library. After preprocessing and quality control, shotgun reads are assembled into contigs, coverage is estimated, and a Hi-C contact matrix is constructed. Three Hi-C-based binners, bin3C, MetaCC, and ImputeCC, independently generate bin sets. METAHICT consolidates them into a non-redundant MAG collection. Intra-contig mapped insert sizes are modeled to select short-insert, shotgun-like Hi-C read pairs, which are combined with shotgun reads for per-bin reassembly. Downstream modules invoke YaHS for Hi-C-guided scaffolding, GTDB-Tk for taxonomic annotation, and geNomad for MGE identification; candidate MGE–host pairs are subsequently evaluated using Hi-C contacts.

METAHICT supports short-read Hi-C libraries paired with short- or long-read shotgun data, and its EM-based read-selection and per-bin reassembly procedures are designed for datasets with paired-end short-read shotgun libraries. The resulting outputs include raw and normalized Hi-C contact matrices, MAG sequences with completeness and contamination estimates, taxonomic assignments, scaffold visualizations, and candidate MGE–host pairs retained after Hi-C contact filtering.

### 2.2 Benchmarking metaHi-C datasets from diverse environments

To evaluate METAHICT across heterogeneous communities and sequencing modalities, we assembled a cross-habitat benchmark comprising seven metaHi-C datasets: human gut [13], sheep gut [24], pig gut [25], cow rumen [26], bovine skin [27], wastewater [10], and hydrothermal mats [28]. Five short-read metaHi-C datasets use short-read shotgun assemblies paired with short-read Hi-C libraries (human gut, pig gut, bovine skin, wastewater, and hydrothermal mats), whereas the sheep-gut and cow-rumen long-read metaHi-C datasets use long-read metagenomic assemblies paired with short-read Hi-C libraries. Dataset characteristics are summarized in Supplementary Table S2.

In the human gut dataset [13], Illumina sequencing produced 125.4 million paired-end shotgun reads (37.9 Gbp) and 85.9 million paired-end Hi-C reads (25.9 Gbp). The pig gut dataset [25], sequenced on an Illumina HiSeq 4000, is of comparable scale, with 86.9 million shotgun read pairs (26.1 Gbp) and 99.3 million Hi-C read pairs (29.8 Gbp). The bovine skin dataset [27] contains 179.5 million shotgun read pairs (53.8 Gbp) and 130.6 million Hi-C read pairs (39.2 Gbp). Among the environmental samples, the wastewater dataset [10] comprises 269.3 million shotgun read pairs (81.3 Gbp) and 95.3 million Hi-C read pairs (28.8 Gbp). For hydrothermal mats [28], we analyzed sample sequenced on an Illumina NovaSeq 6000, yielding 259.3 million shotgun read pairs (77.8 Gbp) and 138.7 million Hi-C read pairs (41.6 Gbp). For the long-read metaHi-C dataset, we used the sheep gut metaHi-C sample [24], which includes PacBio HiFi shotgun sequencing and Illumina HiSeq 2000 Hi-C sequencing. The matched Hi-C library of the sheep gut dataset contained 107.7 million read pairs (32.3 Gbp). The cow rumen long-read metaHi-C dataset [26] includes uncorrected PacBio long-read shotgun sequencing generated on the PacBio RS II and Sequel platforms and short-read Hi-C sequencing. The matched Hi-C library contained 40.9 million read pairs (10.1 Gbp). Restriction enzymes used to construct each Hi-C library are listed in Supplementary Table S2. This collection spans multiple hosts, environments, sequencing technologies, and library sizes, providing a stringent and representative testbed for metaHi-C analysis.

Shotgun reads from the five short-read metaHi-C datasets (human gut, pig gut, bovine skin, wastewater, and hydrothermal mats) were assembled into contigs with MEGAHIT (v1.2.9) [34]. The sheep-gut PacBio HiFi assembly was generated with metaFlye (v2.9) [35] and obtained from Zenodo (see Data Availability). For cow rumen, we used the final assembly reported by Bickhart et al. [26], which was generated from PacBio long-read shotgun data using Canu (v1.6+101 changes, r8513) [36] and subsequently polished twice with Pilon [37]. The published cow-rumen assembly was obtained from Figshare (see Data Availability). Assembly statistics for both datasets are summarized in Supplementary Table S3.

### 2.3 Complementary Hi-C library diagnostics

Because downstream analyses depend on the strength of the Hi-C signal, METAHICT reports two complementary diagnostic indicators. First, the 3D ratio [8], reported by the alignment module, is the fraction of primary read pairs whose mates map unambiguously to different contigs (see Methods, Subsection 4.2). Second, the informative fraction, reported by the reassembly module for short-read datasets, combines the observed inter-contig pairs with the fitted weight of the long-insert component among intra-contig pairs (see Methods, Subsection 4.5). Notably, these quantities are operational diagnostic summaries rather than direct measurements of molecular read origin. The 3D ratio depends on assembly fragmentation and on alignment and filtering choices. The informative fraction additionally assumes that a two-component mixture provides a useful summary of the intra-contig mapped insert-size distribution and operationally treats inter-contig pairs as proximity-ligation products. We report the 3D ratio for all seven datasets and the informative fraction for the five short-read datasets (see Supplementary Table S4). The particularly low 3D ratio for the sheep-gut dataset is consistent with its highly contiguous long-read assembly, in which a larger fraction of ligation products can remain within individual contigs rather than spanning contig boundaries.

### 2.4 Hi-C–informed ensemble binning improves MAG yield and quality across habitats

We evaluated binning performance using CheckM2-estimated completeness and contamination [38], reporting the number of near-complete MAGs (completeness *>*90%, contamination *<*5%) and medium-quality MAGs (completeness *≥*50%, contamination *<*10%) [39]. In head-to-head comparisons against bin3C [17], MetaCC [20], and ImputeCC [30] across seven environments, METAHICT binning module (in short METAHICT in this subsection) consistently produced higher MAG counts across CheckM2 quality thresholds (Fig. 2).

**Fig. 2:**
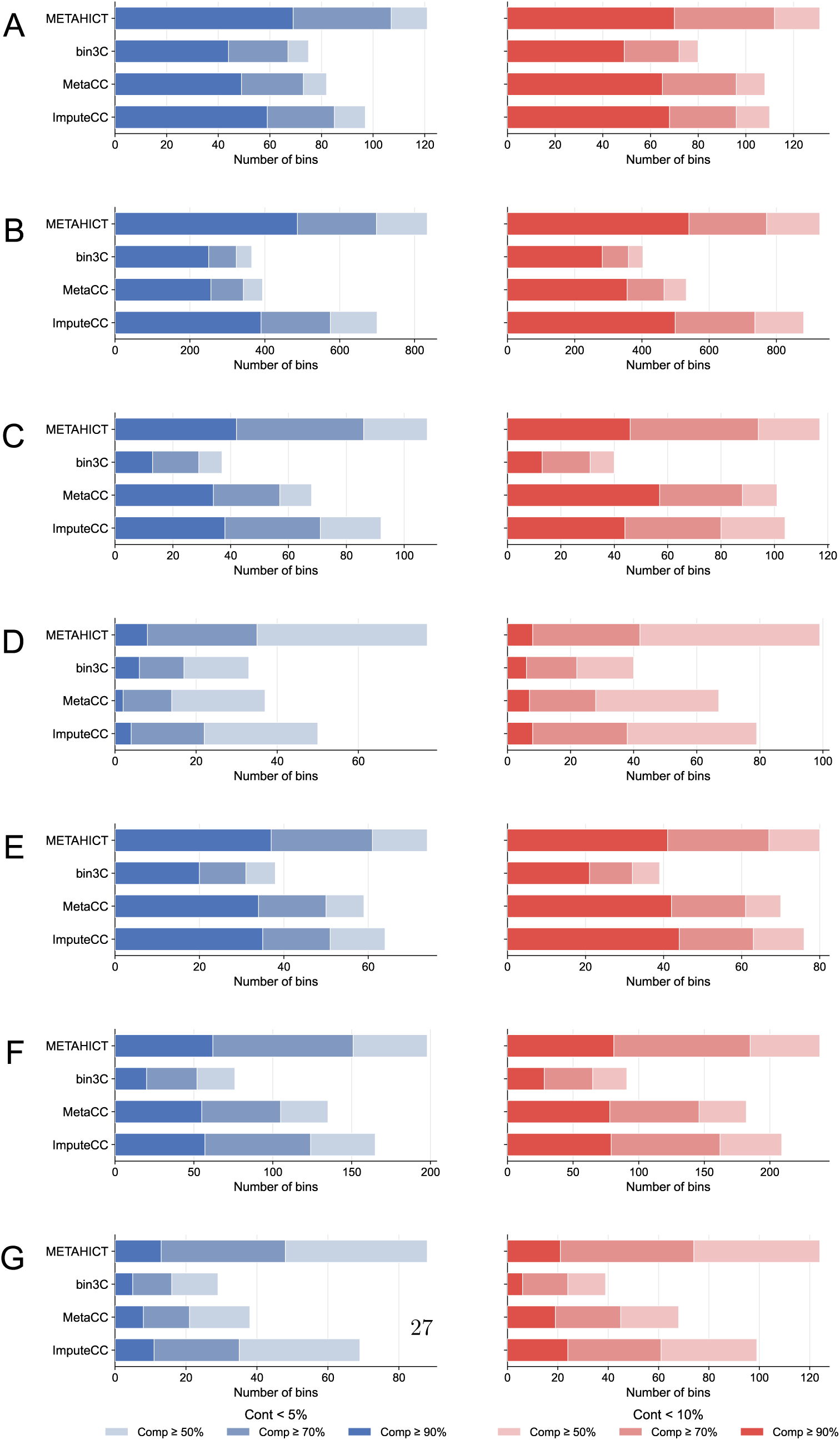
Benchmarking METAHICT binning across seven habitats. Numbers of MAGs meeting quality thresholds (contamination *<*5% or *<*10%; completeness *≥*50%, *≥*70%, or *≥*90%) for METAHICT, bin3C, MetaCC, and ImputeCC in **A** human gut, **B** sheep gut, **C** pig gut, **D** cow rumen, **E** bovine skin, **F** wastewater, and **G** hydrothermal mats. Quality was assessed with CheckM2. Across datasets and thresholds, METAHICT generally recovered more qualifying MAGs than the three individual Hi-C binners.

In the human gut dataset, METAHICT recovered 69 near-complete and 131 medium-quality MAGs, exceeding the next best method (ImputeCC) by 10 (16.9%) and 21 (19.1%), respectively. It also outperformed ImputeCC by retrieving 12.5% more medium-quality bins in the pig gut dataset and 5.7% more near-complete bins in the bovine skin dataset, respectively. In environmental samples, METAHICT assembled 238 (wastewater) and 124 (hydrothermal mats) medium-quality MAGs, improving on ImputeCC by 29 (13.9%) and 25 (25.3%), respectively. In the cow-rumen long-read metaHi-C dataset, METAHICT recovered 8 near-complete and 99 medium-quality MAGs, exceeding the best individual Hi-C binner in each category by 2 and 20 MAGs, respectively. The largest gain was observed in the long-read sheep gut dataset. METAHICT generated 487 near-complete MAGs, surpassing bin3C, MetaCC, and ImputeCC by 237 (94.8%), 231 (90.2%), and 97 (24.9%), respectively. It also produced 929 medium-quality MAGs, exceeding the same methods by 526 (130.5%), 397 (74.6%), and 48 (5.4%), respectively. To quantify the contribution of bin-set consolidation, we compared the METAHICT output with the best-performing individual input binner separately for each dataset and MAG-quality category. Across the seven datasets, consolidation added 2–97 near-complete MAGs, corresponding to gains of 5.7–33.3% relative to the best individual binner, and 4–48 medium-quality MAGs, corresponding to gains of 5.3–25.3%. Dataset-specific absolute and relative gains are reported in Supplementary Table S5. For literature context, MAG counts, numerical quality criteria, taxonomic breadth where available, and binning or refinement workflows reported in the original benchmark studies are summarized in Supplementary Table S6. Since the original studies used different assemblies, workflows, quality-assessment methods, and reporting conventions, these values are presented separately from the controlled benchmarking performed here.

To evaluate the contribution of the METAHICT consolidation procedure, we compared METAHICT with two established bin-ensemble methods using matched inputs. Identical contig-to-bin assignments generated by bin3C, MetaCC, and ImputeCC were supplied to METAHICT, Binning refiner (v1.4.3, default parameters) [40], and DAS Tool (v1.1.7, default parameters) [41]. All outputs were evaluated with CheckM2 using the same quality criteria. METAHICT generally recovered more qualifying MAGs than Binning refiner and DAS Tool across datasets and CheckM2 thresholds (Supplementary Fig. S1). Moreover, we compared the three Hi-C-based binners with the conventional non-Hi-C binners, including MetaBAT2 (v2.18, default parameters) [42] and CONCOCT (v1.1.0, default parameters) [43]. MetaBAT2 and CONCOCT were applied to the same assemblies using contig sequence composition and shotgun-derived coverage information, without Hi-C contact data. Their outputs were evaluated with CheckM2 using the same quality thresholds. Across the seven datasets, the Hi-C-aware methods generally recovered more qualifying MAGs than MetaBAT2 and CONCOCT (Supplementary Fig. S2).

Finally, to assess whether higher counts translate into broader biological coverage, we annotated all medium-quality MAGs using METAHICT annotation module through GTDB-Tk [32] (see Methods, Subsection 4.6). Across datasets, METAHICT generally recovered more distinct GTDB-defined taxa [32] at the species, genus, family, and order levels than the Hi-C–based alternatives (Fig. 3 and Supplementary Fig. S3), indicating that the additional MAGs expand taxonomic breadth rather than duplicating closely related bins. We further quantified redundancy and uniqueness among binners utilizing all-vs-all Mash distances. We calculated distance between medium-quality MAGs produced by METAHICT binning module and other Hi-C–based tools using Mash (v2.3) [44] with the ‘-s 10,000’ parameter. Two MAGs were considered to represent the same underlying genome if their Mash distance was at or below 0.01 [45, 46]. In most comparisons, the majority of MAGs reported by other methods had a match in METAHICT’s set, while METAHICT also contributed a substantial number of unique MAGs not retrieved elsewhere (Fig. 4 and Supplementary Fig. S4). Together, these analyses show that the METAHICT binning module increases the yield of medium-quality MAGs and broadens taxonomic coverage across habitats.

**Fig. 3:**
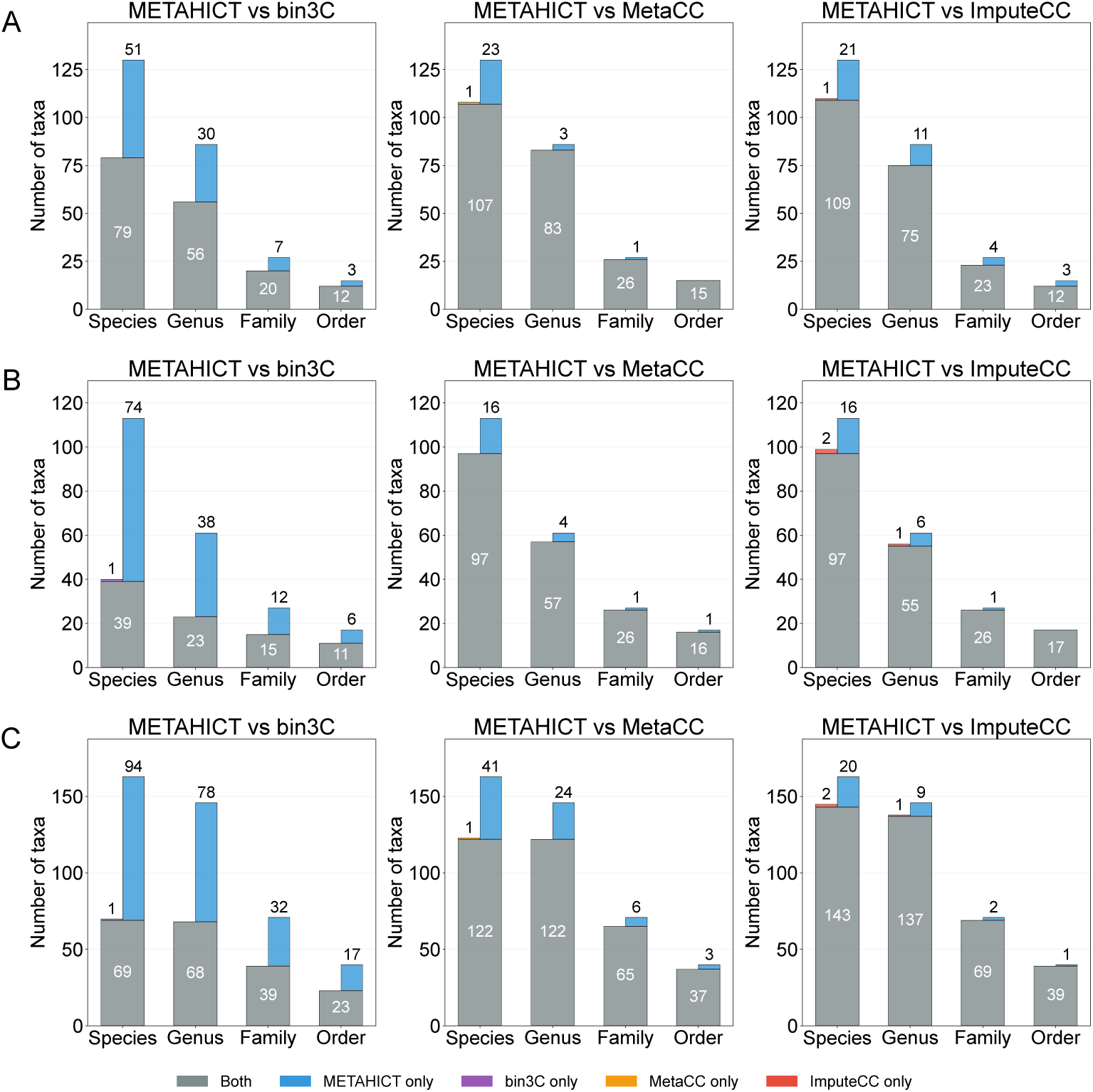
METAHICT binning broadens taxonomic breadth across short- and long-read metaHi-C datasets. Counts of distinct GTDB-defined taxa (species, genus, family, order) from medium-quality MAGs (completeness *≥*50%, contamination *<*10%) in pairwise comparisons between METAHICT binning and each Hi-C–based method (bin3C, MetaCC, ImputeCC). Bars are partitioned into taxa shared by both methods (grey) and taxa unique to one method (colors as indicated). Panels: **A** human gut, **B** pig gut, **C** wastewater. In all datasets, most taxa reported by the alternative methods are also recovered by METAHICT binning, while METAHICT also recovers taxa absent from the comparator outputs.

**Fig. 4:**
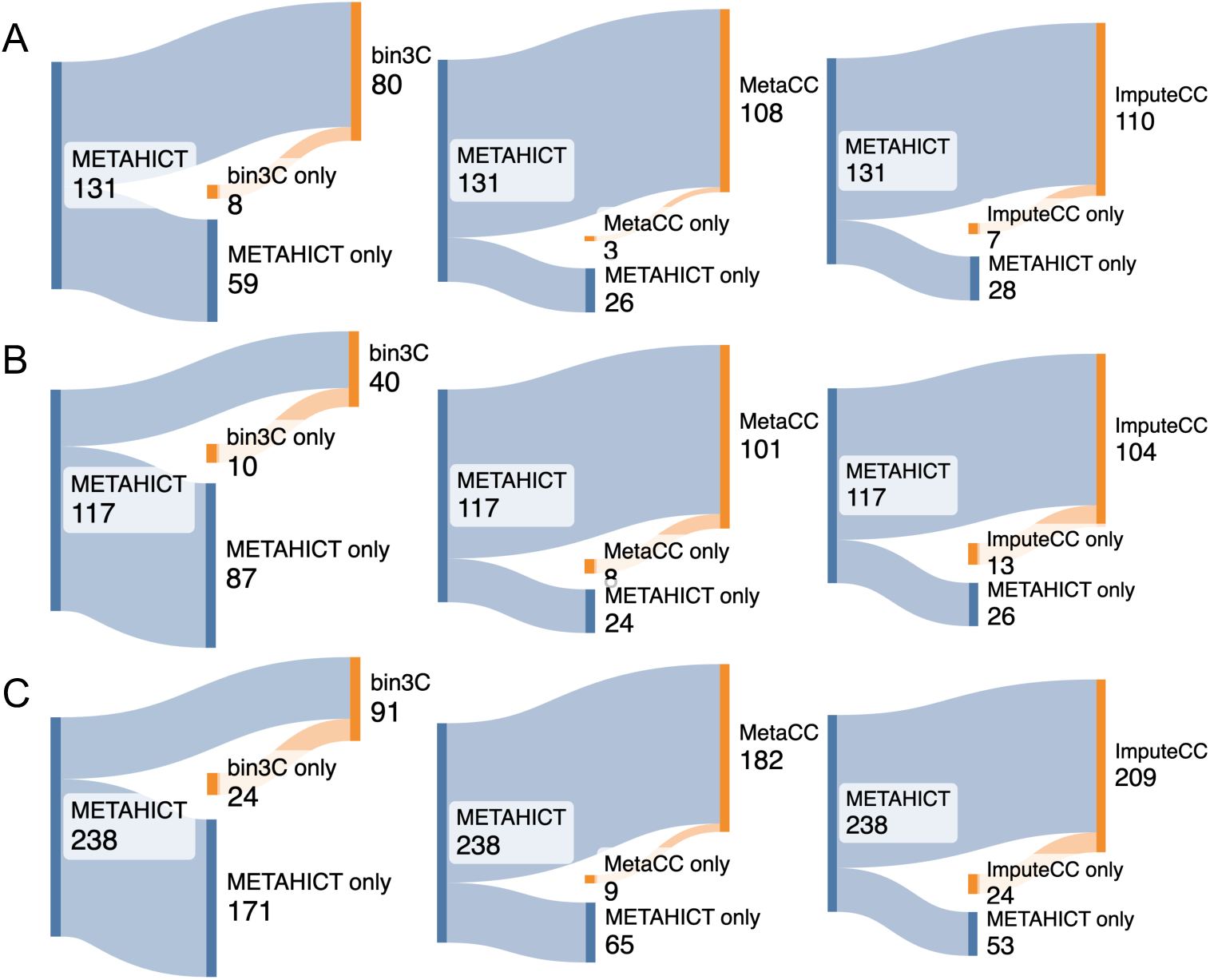
METAHICT binning recovers most overlapping bins and contributes additional unique bins. Sankey diagrams show overlap of medium-quality MAGs (completeness *≥*50%, contamination *<*10%) between METAHICT binning and each Hi-C–based method (bin3C, MetaCC, ImputeCC). Side bars represent all MAGs recovered by each method, while middle bars represent MAGs unique to each method. Panels: **A** human gut, **B** pig gut, **C** wastewater. In all datasets, most bins reported by the alternative methods are also recovered by METAHICT binning, which additionally contributes a substantial number of unique bins.

### 2.5 Per-bin reassembly reduces contamination, with additional gains from EM-selected Hi-C reads

We evaluated the METAHICT reassembly module across the five short-read metaHi-C datasets. The sheep-gut and cow-rumen datasets were excluded because the reassembly module is designed for short-read assemblies. To distinguish the effect of per-bin reassembly from the incremental contribution of EM-selected short-insert read pairs, we evaluated three matched conditions for each starting MAG: the pre-reassembly MAG, reassembly using matched shotgun reads alone (SG-only), and standard METAHICT reassembly using the same shotgun reads plus EM-selected short-insert read pairs from the Hi-C library. The SG-only control was generated by excluding the EM-selected reads while retaining all other reassembly settings. Completeness and contamination were compared between matched MAGs using paired two-sided Wilcoxon signed-rank tests. For each paired contrast, the contamination and completeness tests across the five datasets were adjusted together using the Benjamini–Hochberg procedure [47], resulting in 10 tests per contrast. Adjusted values are reported as *q* values, with *q ≤* 0.05 considered significant (Fig. 5A; Supplementary Table S7).

**Fig. 5:**
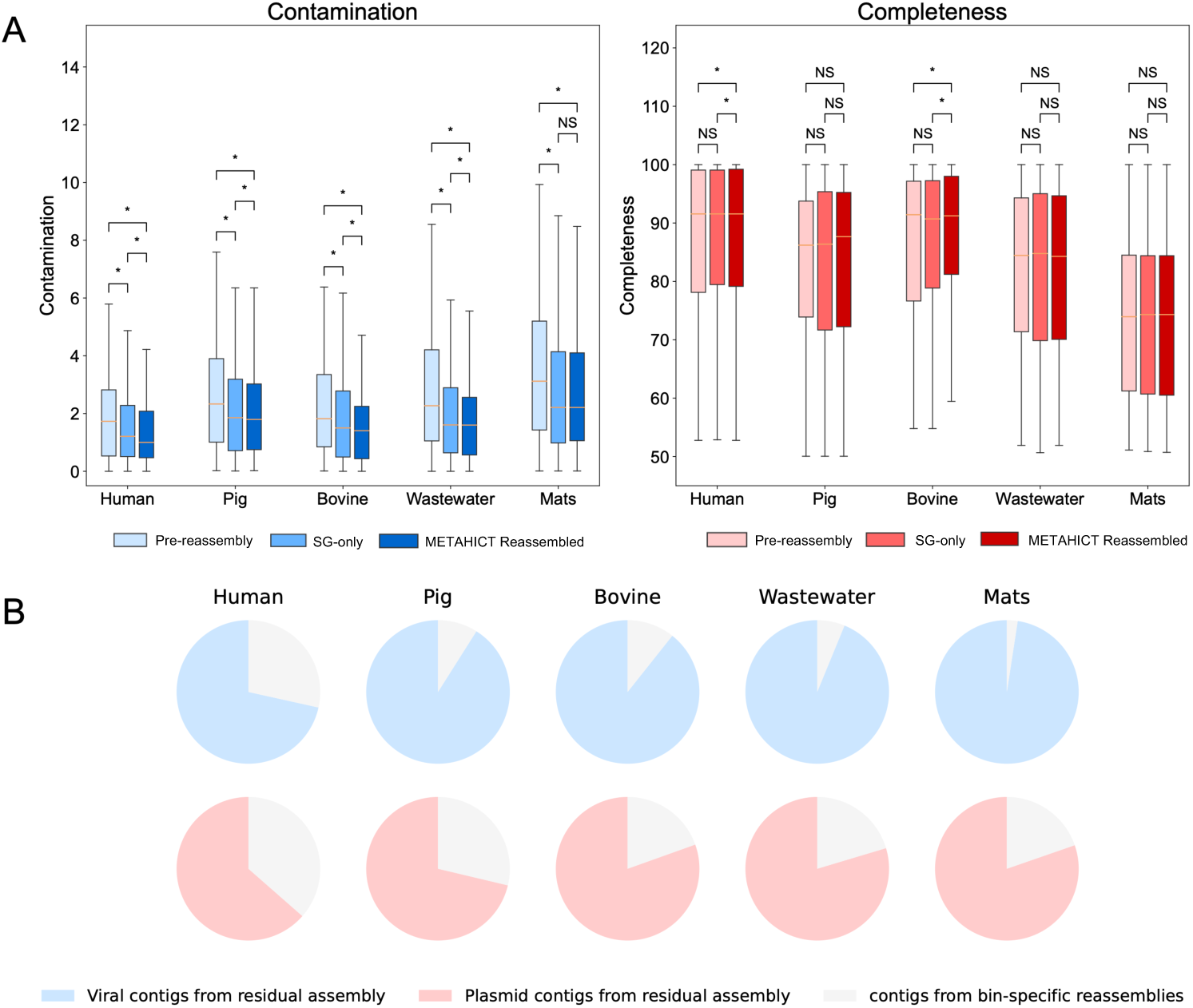
Effects of reassembly, incremental contribution of EM-selected reads, and MGE recovery from residual assemblies across short-read datasets. **A** Paired comparison of CheckM2 contamination (left) and completeness (right) for the pre-reassembly MAGs, reassembly using matched shotgun reads alone (SG-only), and the standard METAHICT reassembly procedure using shotgun reads plus EM-selected short-insert Hi-C reads. Pre-reassembly versus METAHICT reassembly measures the overall effect of the implemented reassembly module and SG-only versus METAHICT reassembly assesses the incremental contribution of the EM-selected reads. From lower to upper, brackets show pre-reassembly versus SG-only, SG-only versus METAHICT reassembly, and pre-reassembly versus METAHICT reassembly. Statistical comparisons used paired two-sided Wilcoxon signed-rank tests, with *p* values for contamination and completeness across the five datasets adjusted together for each paired contrast using the Benjamini–Hochberg procedure. Asterisks indicate *q ≤* 0.05, and NS indicates *q >* 0.05. **B** Residual assemblies contain the majority of detected MGE contigs. For each environment, pie charts show the share of viral (blue) and plasmid (red) contigs identified by geNomad in contigs from bin-specific reassemblies (light wedges) versus contigs assembled from reads not recruited to any bin (darker wedges). Across all five environments, residual assemblies contain more MGE contigs than bin-derived assemblies.

Relative to the pre-reassembly MAGs, standard METAHICT reassembly reduced mean contamination by 0.61–1.02 percentage points across the five datasets. These reductions were significant in all five environments (*q ≤* 3.28 *×* 10*^−^*^9^; Supplementary Table S7). Completeness increased by 0.55 percentage points in human gut (*q* = 1.10*×*10*^−^*^3^) and by 0.92 percentage points in bovine skin (*q* = 4.37*×*10*^−^*^3^), whereas no statistically significant completeness difference was detected in pig gut, wastewater, or hydrothermal mats (*q ≥* 0.815). The SG-only versus standard METAHICT reassembly comparison assessed the incremental contribution of adding the EM-selected read set. Relative to SG-only, inclusion of the EM-selected reads produced further mean contamination reductions of 0.22, 0.13, 0.21, 0.14, and 0.03 percentage points in human gut, pig gut, bovine skin, wastewater, and hydrothermal mats, respectively. These reductions were significant in human gut (*q* = 6.94 *×* 10*^−^*^6^), pig gut (*q* = 4.19 *×* 10*^−^*^3^), bovine skin (*q* = 0.0261), and wastewater (*q* = 6.19 *×* 10*^−^*^5^), but not in hydrothermal mats (*q* = 0.344). Relative to SG-only, completeness increased significantly by 0.19 percentage points in human gut (*q* = 0.0145) and by 0.68 percentage points in bovine skin (*q* = 3.72 *×* 10*^−^*^3^); no significant completeness difference was detected in the other three datasets (*q ≥* 0.106). SG-only reassembly also significantly reduced contamination relative to the pre-reassembly MAGs in all five datasets (*q ≤* 5.43 *×* 10*^−^*^5^), whereas no statistically significant completeness difference was detected in any dataset (*q ≥* 0.0781). Together, these comparisons indicate that shotgun-only per-bin reassembly contributed to the overall contamination reduction, while adding the EM-selected read set provided further significant contamination reductions in four datasets and significant completeness gains in two datasets (Supplementary Fig. S5). Supplementary Table S8 reports N50 and L50 for the MAG collections before and after reassembly. Because reassembly produces a new contig set, these metrics depend on both sequence reconstruction and which contigs are retained and are therefore interpreted only as summaries of the resulting contig-length distributions.

Finally, to avoid losing episomes and other mobile elements that do not co-assemble within chromosomal bins, METAHICT reassembly module also assembles the residual reads that are not recruited to any bin and merges those contigs with bin-specific reassemblies into a single contig set for downstream analyses (see Methods, Subsection 4.5). Applying geNomad (v1.12.0, default parameters) [33] to both sources showed that, across all five environments, residual contigs contain markedly more viral and plasmid sequences than the bin-derived contigs (Fig. 5B). This indicates that much of the MGE sequence content resides outside MAG bins, and relying only on binned contigs for MGE–host linking would therefore miss substantial signal. Assembling and screening the residual read set is thus essential for MGE discovery and downstream MGE–host analysis.

### 2.6 EM-based read selection enriches for short-insert, shotgun-like read pairs from the Hi-C library

METAHICT reuses a subset of intra-contig read pairs from the Hi-C library for per-bin reassembly by modeling their mapped insert-size distribution (see Methods). To evaluate the read-selection procedure using known labels, we used sim3C [48] to generate metagenomic Hi-C read pairs from a mock microbial community with known reference genomes and abundances [49] (see Supplementary Note 1). The simulated reads were aligned to contigs assembled from the original mock-community shotgun library, matching the standard METAHICT workflow. The EM model was fitted using intra-contig read pairs mapped to the default 100 longest assembled contigs, whereas validation was performed using all intra-contig pairs and the sim3C-provided shotgun-derived and Hi-C-derived labels. These labels were not used during model fitting. The statistics of intra-contig Hi-C read pairs and their labels are shown in Supplementary Table S9. Across three simulation seeds, the EM-selected read pairs achieved a mean precision of 84.15%, mean recall of 94.81%, and mean F1 score of 89.16% for shotgun-derived pairs (Supplementary Table S10). On average, the selected set comprised 87.43% of all simulated intra-contig pairs. Thus, the EM procedure enriched for shotgun-derived pairs (Supplementary Fig. S6). We next evaluated the same EM read-selection procedure in the five real short-read metaHi-C datasets. The observed intra-contig mapped insert-size distributions showed a short-insert regime and a long-insert tail, and the fitted EM mixtures placed the read-selection cutoffs near the upper edge of the short-insert regime (Supplementary Fig. S7). The empirical distributions also contained dataset-specific shoulders and tails, consistent with the use of the EM model as a pragmatic read-selection model rather than a complete generative model of all insert sizes. Across the five real datasets, EM-selected reads represented 47.95– 80.50% of all aligned read pairs from the Hi-C library (Supplementary Table S11). Together with the controlled simulation and the shotgun-only reassembly ablation, these results support the enrichment of short-insert, reassembly-compatible reads and their dataset-dependent contribution to reassembly.

### 2.7 Taxonomic composition and species-unclassified MAGs across habitats

Using GTDB-Tk annotations [32] and coverage produced by the annotation and coverage module for medium-quality MAGs (completeness *≥*50%, contamination *<*10%) from the reassembly module for the five short-read datasets and from the binning module for the two long-read datasets, METAHICT generated MAG-resolved taxonomic profiles, species-level richness estimates, and descriptive between-dataset comparisons. Among the seven benchmark datasets, the largest numbers of distinct GTDB-defined species were observed in sheep gut and wastewater (264 and 162, respectively), followed by human gut, pig gut, cow rumen, hydrothermal mats, and bovine skin (130, 113, 87, 43, and 23, respectively). The fraction of bins without a GTDB species label was lowest in human gut (0.8%) and highest in bovine skin (71.2%), showing substantial variation among habitats in the fraction of MAGs assigned at the species level.

Community organization showed clear ecological signatures (Fig. 6A). The three gut habitats as well as the cow rumen environment were dominated by *Bacillota A* and *Bacteroidota*, consistent with fiber- and mucin-associated fermentation [50, 51]; wastewater displayed a mixed profile consistent with heterogeneous inputs and sewer biofilms [52, 53]; hydrothermal mats were enriched for *Campylobacterota* and *Desulfobacterota*, consistent with sulfur cycling [54, 55]; bovine skin featured *Bacteroidota* and *Spirochaetota*, in line with prior reports of *Treponema*-rich communities [56]. Phylum-level Bray-Curtis dissimilarities [29] placed pig gut and cow rumen as the most similar pair, followed by human and sheep gut, whereas pig gut and hydrothermal mats were the most dissimilar (Fig. 6B), showing greater similarity among the gut-associated profiles than between these profiles and the hydrothermal-mat profile.

**Fig. 6:**
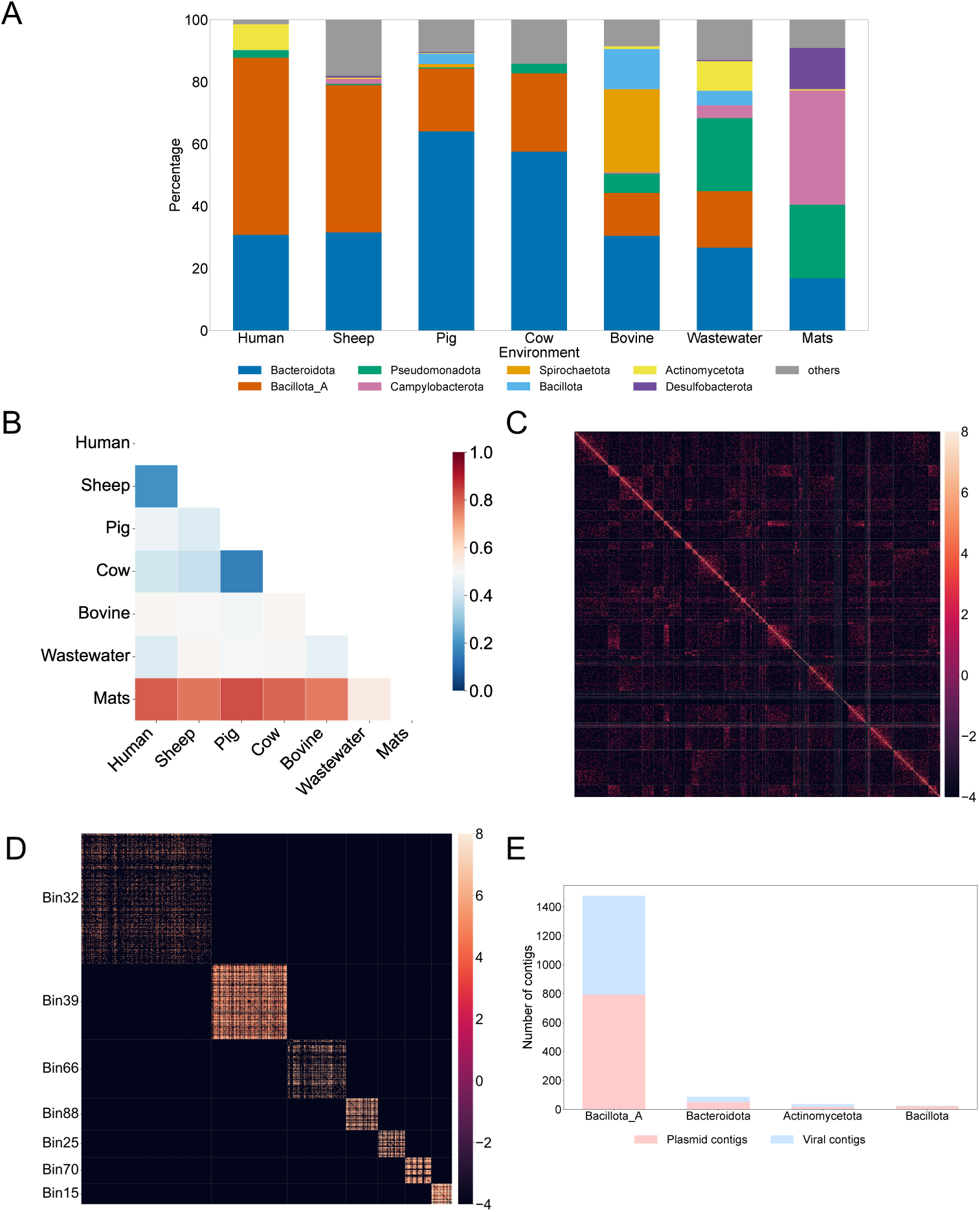
Taxonomic composition, scaffolding, and candidate MGE-MAG associations from METAHICT outputs. **A** Phylum-level community composition (GTDB-Tk annotations) for medium-quality MAGs (completeness *≥*50%, contamination *<*10%) across seven habitats; phyla outside the top eight are grouped as “others”. Gut habitats are dominated by *Bacillota*_*A* and *Bacteroidota*, wastewater shows a mixed profile, hydrothermal mats are enriched for *Campylobacterota* and *Desulfobacterota*, and bovine skin features *Bacteroidota* and *Spirochaetota*. **B** Bray–Curtis dissimilarities among habitats based on phylum-level profiles: pig gut and cow rumen are the most similar pair, followed by human and sheep gut, whereas pig gut and hydrothermal mats are the most dissimilar. **C** Hi-C contact map of scaffolded Bin47 (*Bacteroides vulgatus*), with contacts aggregated between fixed-length genomic intervals. **D** Contig-level within-MAG Hi-C contact matrices for seven *Faecalibacterium* MAGs. Each row and column represents a contig, and the MAGs are ordered by their number of contigs. **E** Counts of viral (blue) and plasmid (red) contigs linked to MAGs from the four most abundant phyla after Hi-C contact filtering. Most retained viral and plasmid contigs were linked to *Bacillota_A* MAGs.

Within selected clades, METAHICT also recovered MAGs with higher-rank taxonomic placements but no species-level GTDB-Tk assignment, here termed species-unclassified MAGs. This designation reflects the taxonomic resolution obtained with GTDB release r220 and does not, on its own, indicate a novel species. In the sheep-gut long-read dataset, METAHICT recovered eight *Erysipelotrichales* MAGs, members of an order associated with host physiology and disease [57]. Four received species-level assignments, whereas one was resolved only to the genus level and three only to the family level. The latter four represent candidates for further taxonomic evaluation.

### 2.8 Scaffolding and MGE analysis in the human gut dataset

In the human gut dataset, the YaHS-based scaffolding module improved contiguity for a dominant *Bacteroides vulgatus* MAG, and the MGE module reported filtered candidate MGE-MAG proximity associations involving MGEs recovered from both bin-derived and residual contigs. We first applied the scaffolding module to Bin47, the most abundant medium-quality MAG. The annotation module assigns it to *Bacteroides vulgatus*, a prevalent gut commensal involved in complex-carbohydrate metabolism [51, 58]. Scaffolding improved contiguity, increasing N50 from 93,828 to 217,798 and decreasing L50 from 19 to 8. The Hi-C contact map of the scaffolded MAG, aggregated into 10-kb genomic intervals (Fig. 6C), showed a clear main diagonal with minimal off-diagonal signal, consistent with structural coherence. Moreover, METAHICT recovered seven *Faecalibacterium* MAGs, a key butyrate-producing genus often depleted in inflammatory bowel disease (IBD) [59, 60], including one species-unclassified MAG (Bin32) without a species-level GTDB-Tk assignment. Bin32 was genetically distinct from the other *Faecalibacterium* MAGs (Mash distance 0.10– 0.20). Contig-level within-MAG contact matrices for the seven *Faecalibacterium* MAGs are shown in Fig. 6D. Scaffolding increased N50 and decreased L50 for all seven *Faecalibacterium* MAGs (Supplementary Table S12).

Applying the MGE module to the human-gut dataset, geNomad (v1.12.0) [33] identified viral and plasmid sequences among the assembled contigs. Contigs annotated as *provirus* were excluded from the standalone viral set. After excluding 152 proviruses, 2,727 viral and 1,817 plasmid contigs remained. Of these, 75% of the viral contigs and 63% of the plasmid contigs originated from the residual assembly, underscoring the contribution of residual-read assembly to MGE recovery. Viral contigs were evaluated with CheckV (v1.0.3, database v1.5) [61], which classified 13 contigs as complete and 20 as high-quality. Candidate virus–host pairs were inferred using the link-level filtering described in Methods, Subsection 4.7. Under the default settings, a candidate pair was retained when it was supported by at least two raw Hi-C read pairs and had a normalized-contact Z-score greater than 0.5. These criteria retained 910 candidate virus–host pairs involving 763 viral contigs and 115 host MAGs. Most retained candidate pairs involved MAGs assigned to *Bacillota A* (Fig. 6E). Within *Bacillota A*, the seven *Faecalibacterium* MAGs differed in their candidate virus–host pairing profiles. The filtered set contained 37 candidate virus–host pairs involving 28 viral contigs, all assigned to *Caudoviricetes* by geNomad. Most viral contigs (22 of 28) were paired with only one of the seven *Faecalibacterium* MAGs, whereas six were paired with two to four MAGs. The *F. hattorii* MAG had the largest candidate set, with 13 pairs, and 10 of these viral contigs were paired only with *F. hattorii* among the seven MAGs examined. In contrast, all three viral contigs paired with the *F. prausnitzii* MAG also formed candidate pairs with at least one other *Faecalibacterium* MAG. Per-pair raw-contact support, normalized-contact scores, Z-scores, CheckV quality categories, and viral and host taxonomic annotations are provided in Supplementary Data 1. Together, these filtered pairs show variation in candidate virus–host pairing profiles among the seven *Faecalibacterium* MAGs, with some viral contigs paired with a single MAG and others with multiple MAGs within this comparison. These candidate pairings provide a focused set for orthogonal validation. Finally, sequence-topology assessment of the human-gut dataset identified 50 MGE contigs annotated by geNomad with DTR or ITR and 45 MGE contigs with ccfind terminal-overlap evidence consistent with putative circularity (Supplementary Table S13). Forty-two contigs had both types of evidence. Ccfind also identified terminal-overlap evidence in 96 non-MGE contigs. Corresponding results for all seven datasets are provided in Supplementary Table S13.

### 2.9 Runtime and computational cost

Module-level runtime and peak memory usage across the seven benchmark datasets are summarized in Supplementary Table S14. Measurements were obtained on an AMD EPYC 9654 96-core processor with 1,536 GB of memory, using up to 80 threads per workflow run. Binning required 0.59–29.82 h and 13.40–133.74 GB of peak memory. Across the five short-read datasets, reassembly required 5.01–11.97 h and 30.31–72.51 GB of peak memory.

## 3 Discussion

METAHICT provides an integrated metaHi-C workflow from raw reads to genome-resolved outputs, with separately executable modules, documented defaults, and configurable analytical and resource settings. Selected parameters can be adjusted to account for differences in data quality, sequencing depth, and available computational resources. Across seven habitats, it increased recovery of near-complete and medium-quality MAGs over established Hi-C baselines, while per-bin reassembly reduced contamination with little impact on completeness. Habitat-level patterns derived from length-weighted profiles aligned with ecological expectations, and focused cases highlighted biologically informative genomes and interactions. The two long-read benchmarks extend the evaluation of METAHICT binning and consolidation beyond short-read assemblies. In both the sheep-gut and cow-rumen datasets, METAHICT recovered more near-complete and medium-quality MAGs than the best-performing individual Hi-C binner. Both long-read assemblies had higher mean contig lengths than the five short-read assemblies (Supplementary Table S3). Differences in MAG recovery may reflect multiple biological and technical factors, including community composition, sequencing depth, and assembly characteristics. The EM-based per-bin reassembly module is designed and evaluated for short-read assemblies.

Two workflow-level procedures contributed to these gains. First, treating Hi-C–informed binning as an ensemble task and consolidating multiple bin sets into a non-redundant collection avoids dependence on any single method and improves yield and breadth. Second, reusing intra-contig signal by selecting short-insert, shotgun-like read pairs from Hi-C libraries with an EM model on mapped insert sizes and supplying them to reassembly offers a practical route to cleaner drafts. The sim3C validation showed that the EM procedure enriches for shotgun-derived read pairs while reducing the inclusion of longer-insert Hi-C-derived pairs. Because short-range Hi-C-derived pairs can overlap shotgun-derived pairs in mapped insert size, we interpret this step as an enrichment procedure for short-insert, reassembly-compatible read pairs rather than as an exact molecular-origin classifier. Alignment-derived indicators (3D ratio [8]) and the model-based informative fraction provide complementary diagnostics of long-range signal. METAHICT also connects the binning and reassembly outputs to established downstream tools: YaHS supports Hi-C-guided scaffolding, GTDB-Tk provides taxonomic assignment, and geNomad identifies candidate MGE sequences. Hi-C contacts are then used to report candidate MGE–host pairs for further evaluation. In practice, reassembly is compute-intensive and benefits from adequate coverage; METAHICT therefore supports selective use (e.g., prioritizing higher-contamination or high-value MAGs) to match available resources. Where resources are constrained, users may skip this step and proceed with the consolidated bins from the METAHICT binning module for downstream analyses, although our results show that reassembly significantly reduced contamination of the retrieved MAGs.

In the MGE module, Hi-C contact enrichment is used to prioritize candidate MGE– host pairs. The raw-contact and normalized-contact thresholds apply reproducible empirical filters, but they do not provide formal significance testing or false-discovery-rate control. Candidate MGE–host pairs are therefore regarded as hypotheses for validation rather than as confirmed infections or exclusive host assignments. Evaluation of individual pairs will require independent evidence, such as longitudinal recurrence or experimental validation. Future work should develop statistically calibrated null models and systematically benchmark them across community types, sequencing depths, and Hi-C protocols. METAHICT reports geNomad terminal-repeat annotations for MGE contigs and ccfind terminal-overlap evidence for all assembled contigs. These annotations can prioritize sequences for further evaluation, although neither terminal repeats nor terminal overlaps alone establish molecular circularity or demonstrate that a MAG represents a closed chromosome. They complement CheckM2-based estimates of genome completeness and contamination by describing a distinct aspect of assembly structure.

Looking ahead, useful extensions include strain-aware binning and phasing, longitudinal tracking of MAGs and candidate MGE–host associations, and data-driven thresholding for spurious-contact removal. These areas are active and evolving, so we do not prescribe them here; instead, METAHICT is structured to accommodate such advances as they mature. In summary, by combining ensemble binning, EM-guided read reuse, and integrated downstream analysis within a reproducible framework, METAHICT provides standardized MAG, taxonomic, contact-map, scaffolding, and MGE–host outputs for comparative and hypothesis-driven microbiome studies.

## 4 Methods

### 4.1 Implementation

METAHICT is implemented as a Nextflow DSL2 workflow [22], which coordinates module execution and software environments. The workflow can be run end to end or initiated from a supported module when the required upstream inputs are available. An editable YAML configuration specifies analytical and resource settings. Software dependencies are managed through versioned conda environments and checksum-verified lock files. For each run, METAHICT records the configuration-file path and checksum, software versions, process logs, and standard Nextflow execution reports. The analytical settings used in this study are summarized in Supplementary Table S1, and the complete parameter and default-value reference is available in the software documentation.

### 4.2 Processing Raw Data

METAHICT provides an integrated workflow for processing raw metaHi-C sequencing reads, comprising four modules: Preprocessing, Assembly, Alignment, and Coverage.

#### Preprocessing module

Read cleaning is required before aligning Hi-C pairs because adaptor/linker sequences and low–quality bases confound downstream analyses. METAHICT cleans shotgun and Hi-C libraries with BBduk (BBTools v39.10) [62]. The preprocessing module exposes BBduk parameters controlling adaptor removal (– ktrim, –k, –mink, and –hdist), end-quality trimming (–qtrim, –trimq, and –ftm), fixed 5’ trimming (–ftl), and minimum post-trim read length (–minlen). By default, the benchmark analyses used adaptor removal with ktrim=r, k=23, mink=11, hdist=1, tpe, and tbo; end-quality trimming with qtrim=r, trimq=10, and ftm=5; fixed 5’ hard trimming with ftl=10; and a minimum post-trim read length of 50 bp. Subsequently, FastQC (v0.12.1) [63] is employed to assess read quality and evaluate the effectiveness of preprocessing.

#### Assembly module

METAHICT assembles short-read shotgun libraries with MEGAHIT [34] using a broad k–mer sweep (v1.2.9, default options: -k-min 21, -k-max 141, -k-step 12, -merge-level 20,0.95). metaSPAdes (v4.0.0) [64] is provided as an alternative. For long-read shotgun libraries, assemblies are generated with metaFlye (v2.9.5) [35].

#### Alignment module

METAHICT aligns preprocessed paired-end Hi-C reads to assembled contigs using BWA-MEM (v0.7.18) [65]. The BWA command string is user-adjustable, with -5SP used by default because it disables pairing mode and retains the alignment with the lowest read coordinate as primary [66]. Post-alignment filtering with SAMtools (v1.21) [67] removes unmapped reads and excludes secondary, supplementary, and low-quality alignments. The filtering parameters are also exposed, including the SAM flag filter, minimum mapping quality, and minimum nucleotide match length. By default, the analyses in this study removed secondary and supplementary alignments using -F 0x900 and retained alignments with nucleotide match length ≥ 30 and mapping quality ≥ 30. METAHICT also reports a library-level 3D ratio [8] to quantify enrichment for between-contig contacts, defined as the ratio of primary read pairs whose mates map to two different contigs to the total number of primary read pairs.

#### Coverage module

METAHICT computes the coverage of contigs using the MetaBAT2 (v2.18) [42] script ‘jgi_summarize_bam_contig_depths’ with default parameters.

### 4.3 Generating raw and normalized Hi-C contacts

#### Contact module

The assembled contigs, Hi-C read alignment results, and restriction enzyme(s) information serve as inputs to the contact module. Short contigs with limited Hi-C signals typically exhibit higher variance, reducing stability in downstream analyses [17, 68]. METAHICT therefore applies user-adjustable filtering criteria for minimum contig length and minimum number of across-contig Hi-C contacts. By default, the analyses in this study excluded contigs shorter than 1,000 bp and contigs containing fewer than one across-contig Hi-C contacts. The raw Hi-C contact matrix is then constructed from paired-end Hi-C read alignments, with diagonal entries representing intra-contig contacts and off-diagonal entries representing inter-contig contacts.

Since raw metaHi-C contacts contain systematic biases and noise that compromise the reliability of microbial interaction networks, normalization is essential prior to downstream analyses [14]. METAHICT uses a two-step normalization procedure: bias correction followed by spurious-contact filtering. In this study, raw Hi-C contact matrices were normalized using NormCC (v1.2.0) [20]. The default lowest 5% of positive normalized contact values were removed as potential spurious interactions [20, 68]. Alternative normalization methods implemented in METAHICT, including bin3C [17], HiCzin [68], and MetaTOR [18], are summarized in Supplementary Note 2. The workflow configuration allows users to select the contact-normalization method and adjust the corresponding filtering and normalization settings.

### 4.4 Hi-C-informed ensemble binning and MAG refinement

#### Binning module

The binning module of METAHICT integrates multiple Hi-C-based binning approaches to produce a refined set of metagenome-assembled genomes (MAGs). Specifically, we applied three Hi-C-informed binning methods, including bin3C (v0.1.1a) [17], MetaCC (v1.2.0) [20], and ImputeCC (v1.0.0) [30], using their default parameter settings. All three supported Hi-C-based binners were run within the binning module while their method-specific parameters could be adjusted through the workflow configuration. The resulting bin assignments were combined in the ensemble-refinement procedure described below. Because bin3C was originally implemented in Python 2, we ported it to Python 3 for use in METAHICT.

The outputs of these individual binning methods are systematically combined using a hybrid refinement strategy inspired by [69]. First, hybrid candidate bin sets are generated from the bin3C, MetaCC, and ImputeCC assignments using an adapted implementation of Binning refiner [40], producing pairwise and three-way intersections of the original bin sets. This process carefully resolves contigs assigned inconsistently across different binning tools, resulting in hybrid bins that integrate complementary strengths from multiple predictions. Subsequently, bins from the three original and four hybrid bin sets were evaluated with CheckM2 (v1.1.0, default parameters) [38]. Bins with estimated completeness of at least 50% and contamination of at most 10% were retained as candidates for consolidation. Within each duplicate group, the representative bin is selected by maximizing

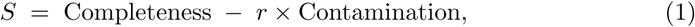

where *r* is a user-adjustable contamination penalty weight that is set to its default value of 5. Ties are resolved by preferring higher completeness, then lower contamination, and finally larger assembly size. After selecting one representative per group, the consolidated set is formed. Finally, any contig appearing in more than one retained bin is assigned uniquely to the highest-scoring bin under the same criterion, and CheckM2 is rerun on the resulting non-redundant collection to refresh quality metrics.

### 4.5 EM-based selection of short-insert, shotgun-like intra-contig Hi-C read pairs for reassembly

#### Reassembly module

Hi-C libraries can contain short-insert read pairs that do not provide useful long-range proximity information and instead resemble conventional whole genome shotgun read pairs because Hi-C enrichment is incomplete [48, 70]. Although such pairs are typically not used for inter-contig contact analysis, they can still provide short-read sequence information for MAG-specific reassembly. METAHICT therefore selects short-insert, shotgun-like intra-contig Hi-C read pairs and reuses them for per-bin reassembly. All Hi-C read pairs mapped to different contigs are treated as proximity-ligation-derived products and are not reused for reassembly. For read pairs whose two mates map to the same contig, METAHICT extracts the mapped coordinates from the BAM file and computes the intra-contig mapped insert size,

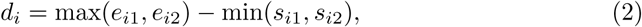

where *s_i_*_1_ and *e_i_*_1_ denote the start and end coordinates of mate 1, and *s_i_*_2_ and *e_i_*_2_ denote the start and end coordinates of mate 2. We use the term mapped insert size to distinguish this coordinate span from the inner gap between mates. The empirical distribution of *d_i_* is modeled as containing a short-insert component enriched for shotgun-like read pairs and a longer-insert component enriched for proximity-ligation-like products. To separate these regimes, METAHICT fits a two-component Gaussian mixture to the intra-contig mapped insert sizes,

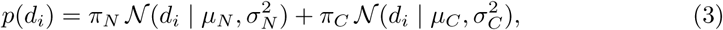

where component *N* denotes the short-insert, shotgun-like component and component *C* denotes the longer-insert, proximity-ligation-like component. Parameters are estimated by expectation–maximization (EM) [23]. METAHICT uses intra-contig read pairs mapped to the *k* (default: 100) longest assembled contigs. This top-contig fitting strategy focuses EM estimation on contigs with sufficient genomic span to represent both the short-insert regime and the longer-insert intra-contig Hi-C tail, while avoiding the strong truncation of insert-size distributions on short contigs. The *N* and *C* components were initialized using the lower 80% and upper 20% of mapped insert sizes, respectively. EM fitting terminated when the absolute change in log-likelihood was less than 0.01 or when 100 iterations had been completed. Following convergence, the estimated cutoff was applied to all filtered intra-contig Hi-C read pairs. A read pair was selected as short-insert and shotgun-like when

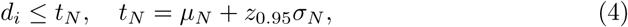

where *z*_0.95_ is the 95th percentile of the standard normal distribution. The 95th-percentile cutoff retains the main short-insert regime while excluding longer-insert intra-contig read pairs. All inter-contig Hi-C read pairs are treated as proximity-ligation products and are not reused for reassembly. The EM-selected intra-contig pairs were retained as a read set enriched for short-insert, shotgun-like pairs and used in subsequent per-bin reassembly.

The fitted *C*-component weight is reported as a model-based indicator of the long-insert, proximity-ligation-like component among intra-contig pairs. Because short-range Hi-C products can overlap the short-insert component, this weight should not be interpreted as the total molecular fraction of proximity-ligation-derived intra-contig read pairs. An operational informative-fraction diagnostic is estimated as

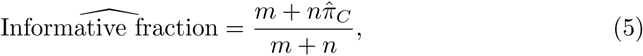

where *m* is the number of inter-contig pairs, *n* is the number of intra-contig pairs considered by the model, and *π*^*_C_* is the fitted long-insert component weight. This estimate is a diagnostic summary rather than ground truth and is interpreted alongside the 3D ratio [8].

Read identifiers for EM-selected short-insert read pairs are exported and merged with the matched shotgun reads. METAHICT then performs per-bin reassembly using both sources, employing the reassembly framework from [69] to improve MAG quality. Further methodological details are provided in Supplementary Note 3. Selected parameter settings used in this study are summarized in Supplementary Table S1. After per-bin read recruitment and reassembly, residual reads that are not assigned to any bin are pooled and assembled de novo with MEGAHIT [34] (v1.2.9; default options: -k-min 21, -k-max 141, -k-step 12, -merge-level 20,0.95). This residual assembly preserves episomes and other mobile elements that do not co-assemble within chromosomal bins and would otherwise be lost. Finally, METAHICT merges contigs from (i) the bin-specific reassemblies and (ii) the residual assembly into a single contig set for downstream analyses. The reassembly module of METAHICT is intended for short-read metaHi-C datasets and is not applied to long-read datasets to avoid mixing assembly paradigms; a comparable hybrid refinement for long-read data is outside the scope of this study.

### 4.6 MAG scaffolding and annotation

#### Scaffolding module

METAHICT scaffolds and visualizes MAGs using Hi-C contacts. Given a MAG’s contigs and the corresponding Hi-C alignments, METAHICT first runs YaHS (v1.2.2; default parameters) to infer scaffold order and orientation from contact patterns [31]. After scaffolding, a Hi-C contact heatmap is generated for each MAG from the scaffolded assembly and its corresponding contact matrix. Contacts are aggregated within fixed-length genomic bins, with a bin size of 10 kb used in this study. Each heatmap cell represents the contact signal between a pair of genomic bins. Continuity along the main diagonal provides a visual measure of scaffold structure, whereas discontinuities or unexpected off-diagonal signals may indicate potential misjoins or gaps. YaHS settings and heatmap resolution are specified through the workflow configuration.

#### Annotation module

METAHICT provides an annotation module with GTDB-Tk (v2.4.0) [32] using the ‘classify_wf’ workflow and GTDB database (release r220), providing standardized, rank-consistent taxonomic assignments that enhance biological interpretability.

### 4.7 Detecting and analyzing mobile genetic elements

#### MGE module

METAHICT identifies mobile genetic elements (MGEs), including viral and plasmid contigs, and reports candidate MGE–MAG pairs. MGE identification and annotation are performed with geNomad (v1.12.0; default parameters) [33]. Contigs or regions annotated by geNomad as proviruses are excluded from the standalone MGE set to avoid conflating host-associated proviral sequence with independently assembled MGEs. Candidate-pair inference uses both the raw and normalized Hi-C contact matrices generated by the METAHICT contact module. Candidate MGE– MAG pairs are enumerated from normalized contacts between each MGE contig and the non-MGE contigs assigned to a MAG. For every candidate pair, METAHICT reports the number of raw Hi-C read pairs linking the MGE contig to the MAG and the aggregated normalized-contact score across the corresponding MGE–MAG contacts. Let *S_vm_*denote the aggregated normalized-contact score between MGE *v* and MAG *m*. Within each sample *s*, this score is standardized as 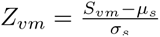, where *µ_s_* and *σ_s_* are the mean and standard deviation, respectively, of the positive aggregated normalized-contact scores in that sample. Under the default settings, a candidate MGE–MAG pair is retained when it is supported by at least two raw Hi-C read pairs and has a normalized-contact Z-score greater than 0.5. The minimum raw-contact requirement follows previous Hi-C-based MGE-linking studies [6, 9, 71], whereas the default Z-score threshold was selected based on the synthetic-community benchmarking reported by Shatadru et al. [72]. These thresholds constitute an empirical pair-level filtering strategy and users can modify the filtering strategy and both thresholds through the module options.

The MGE module also reports contig-level sequence-topology annotations. For viral and plasmid contigs identified by geNomad, METAHICT records whether geNomad reports direct terminal repeats (DTR), inverted terminal repeats (ITR), or no detected terminal repeats. These categories are treated as terminal-repeat annotations rather than as definitive evidence of molecular circularity. In parallel, ccfind (v1.4.7) [73] is applied to all assembled contigs. The default thresholds are a terminal-fragment size of 500 bp, a minimum aligned length of 50 bp, and a minimum sequence identity of 94%; all three thresholds are user-configurable. Contigs meeting these criteria are annotated as having terminal-overlap evidence consistent with putative circularity.

## Declarations

## Acknowledgements

Y.D. is partially supported by the University of Texas Systems STARs Program.

## Consent for publication

All authors have approved the manuscript for submission.

## Data Availability

Raw sequencing data for the benchmark datasets are publicly available through the NCBI Sequence Read Archive (SRA) and the European Nucleotide Archive (ENA). The human-gut shotgun reads are available under SRR6131123, and the corresponding Hi-C reads are available under SRR6131122 and SRR6131124. The pig-gut shotgun reads are available under ERR7197595–ERR7197599, and the corresponding Hi-C reads are available under ERR7197655. The bovine-skin shotgun and Hi-C reads are available under SRR13765540 and SRR13765539, respectively. The wastewater shotgun and Hi-C reads are available under SRR8239393 and SRR8239392, respectively. The hydrothermal-mat shotgun and Hi-C reads are available under SRR21545383 and SRR22355230, respectively. The sheep-gut sequencing data are available under BioProject PRJNA595610. The PacBio HiFi assembly used in this study is available at https://doi.org/10.5281/zenodo.5228989 under the filename flye.v29.sheep_gut.hifi.250g.fasta.gz. The cow-rumen sequencing data are available under BioProject PRJNA507739, and the final polished assembly used in this study is available at https://figshare.com/articles/dataset/usda_pacbio_second_pilon_indelsonly_fa_gz/8323154. The mock-community shotgun data used for the controlled sim3C validation are available from ENA under project PRJEB52977. The remaining data supporting the findings of this study are provided in the Article, Supplementary Information, and Supplementary Data. There is no restriction on data availability. Source data are provided with this paper.

## Code Availability

The METAHICT software is publicly available at https://github.com/dyxstat/METAHICT under the GNU General Public License version 3. The METAHICT code used in this work is also archived on Zenodo under https://doi.org/10.5281/zenodo.22310633. Default values for configurable analytical parameters and settings are provided in the YAML configuration file at https://github.com/dyxstat/METAHICT/blob/main/nextflow/assets/metahict_configuration.yaml. A compact example dataset for testing the workflow is included in the repository at https://github.com/dyxstat/METAHICT/tree/main/example_dataset. Scripts used in this study to process intermediate data and generate the figures are available at https://github.com/dyxstat/Reproduce_METAHICT.

## Ethics approval and consent to participate

Not applicable.

