## Supplementary Note; Supplementary Figure; Supplementary Table for "METAHICT enables comprehensive genome-resolved microbiome analysis with metagenomic Hi-C"

### Supplementary Notes

#### Supplementary Note 1: Validation and model-fit assessment of EM-based read selection

To validate the EM-based read-selection step with known labels, we used sim3C v0.2 [1] to generate metagenomic Hi-C reads from mock-community reference genomes with known species abundances. The mock-community shotgun data were downloaded from the European Nucleotide Archive under project ID PRJEB52977 [2]. The mock community comprises 71 strains representing 69 distinct species and underwent comprehensive whole genome shotgun sequencing using the Illumina HiSeq 3000. After excluding incomplete reference genomes, we retained reference genomes representing 66 species for sim3C read generation. Species abundances were obtained from the supplementary data of Meslier et al [2]. The accession number of the shotgun library is ERR9765746 with 6.1 Gbp shotgun reads. Hi-C reads were simulated using the command `-n 10000000 -l 150 -e MluCI -e Sau3AI -m hic -insert-sd 20 -insert-mean 350 -insert-min 150 -linear -simple-reads` [3, 4]. To account for stochastic variation in the simulation, we repeated the analysis with three random seeds: `-seed 101`, `-seed 102`, and `-seed 103`. To mimic the actual METAHICT workflow, the simulated Hi-C reads were aligned to contigs assembled from the original mock-community shotgun library. After assembly, a total of 15,214 contigs longer than 1,000 bp were obtained, with an average length of 10,535.18 bp. The same BWA-MEM and post-alignment filtering settings used by METAHICT were applied. Intra-contig mapped insert sizes were then extracted from the resulting alignments, and the METAHICT EM procedure was applied. The sim3C-provided shotgun/Hi-C labels were not used during EM fitting and were used only for post hoc evaluation. Shotgun-derived intra-contig read pairs were treated as the positive class and Hi-C-derived intra-contig read pairs as the negative class. Read-selection performance was summarized using precision, recall, and F1 score, with shotgun-derived intra-contig read pairs treated as the positive class for this controlled evaluation. Because short-range Hi-C-derived pairs can overlap shotgun-derived pairs in mapped insert size, these metrics assess enrichment for shotgun-derived reads and should not be interpreted as evidence of exact molecular-origin classification.

For each real short-read metaHi-C dataset, EM fitting used intra-contig read pairs mapped to the 100 longest assembled contigs, matching the default METAHICT workflow. The fitted cutoff was subsequently applied to all intra-contig Hi-C read pairs. For visualization, the mapped insert-size distributions were downsampled where necessary to reduce plotting density; this visualization-only downsampling was not used for EM fitting or for calculation of the selected-read fractions. For each dataset, we report the number of aligned Hi-C read pairs, the number of read pairs used for model fitting, the number of EM-selected read pairs, and the selected fractions relative to all aligned Hi-C read pairs.

#### Supplementary Note 2: Normalization strategies implemented in METAHICT for correcting systematic biases

##### bin3C

bin3C [3] applies a two-step normalization approach: restriction site normalization followed by bistochastic matrix balancing.

The first step corrects for biases introduced by restriction enzyme digestion, as Hi-C contacts originate from ligation at enzyme cut sites. Given a raw Hi-C contact count $H_{ij}$ between contigs $i$ and $j$, the adjusted interaction count $H'_{ij}$ is computed as:

$$H'_{ij} = \frac{H_{ij}}{s_i \cdot s_j}, \tag{S1}$$

where $s_i$ and $s_j$ denote the number of restriction sites on contigs $i$ and $j$, respectively.

Following restriction site normalization, bin3C applies bistochastic matrix balancing using the Knight-Ruiz algorithm [5]. This ensures that each row and column of the normalized Hi-C contact matrix sums to a uniform value. The balancing process iteratively computes a diagonal scaling matrix $D$ such that:

$$DH'D = \tilde{H}, \tag{S2}$$

where $\tilde{H}$ is the balanced contact matrix. The diagonal matrix $D$ is iteratively adjusted to satisfy:

$$\sum_j \tilde{H}_{ij} = 1, \quad \sum_i \tilde{H}_{ij} = 1. \tag{S3}$$

##### MetaTOR

MetaTOR [6] constructs a normalized Hi-C contact network to correct for contact-frequency biases. Intercontig Hi-C contact counts $x_{ij}$ are normalized by the geometric mean of the corresponding intra-contig contacts, yielding a score:

$$S_{ij} = \frac{H_{ij}}{\sqrt{x_{ii} \cdot x_{jj}}}, \tag{S4}$$

where $x_{ii}$ and $x_{jj}$ denote the numbers of intra-contig contacts within contigs $i$ and $j$, respectively.

##### HiCzin

HiCzin (Unlabeled Version) [7] models Hi-C contact counts using a negative binomial distribution:

$$H_{ij} \sim \mathrm{NB}(\mu_{ij}, \theta), \tag{S5}$$

where $\mu_{ij}$ is the expected contact frequency, and $\theta$ is the dispersion parameter.

To correct for bias factors, $\mu_{ij}$ is modeled as:

$$\log(\mu_{ij}) = \beta_0 + \beta_s \log(s_i s_j) + \beta_l \log(l_i l_j) + \beta_c \log(c_i c_j), \tag{S6}$$

where $s_i, l_i, c_i$ denote the restriction sites, contig length, and coverage of contig $i$, respectively.

Normalized Hi-C contacts are computed as residuals:

$$e_{ij} = \frac{H_{ij}}{\widehat{\mu_{ij}}}. \tag{S7}$$

##### NormCC

NormCC [8] models the total Hi-C signal per contig to eliminate biases associated with restriction sites, contig length, and sequencing coverage. The Hi-C signal for contig $i$, defined as the total number of proximity ligation events across all contigs, is:

$$M_i = \sum_{k \neq i} H_{ik}. \tag{S8}$$

NormCC models $M_i$ using a negative binomial distribution:

$$M_i \sim \mathrm{NB}(\mu_i, \theta), \tag{S9}$$

where $\mu_i$ is linked to explicit bias factors via:

$$\log(\mu_i) = \beta_0 + \beta_s \log(s_i) + \beta_l \log(l_i) + \beta_c \log(c_i). \tag{S10}$$

Since $c_i$ is often unknown, it is estimated from within-contig Hi-C contacts:

$$c_i \approx \left( \frac{N_i}{s_i^{\gamma_s} l_i^{\gamma_l}} \right)^{-\gamma_c}, \tag{S11}$$

where $N_i = H_{ii}$ is the within-contig Hi-C contact count.

The final normalized Hi-C contacts between contigs $i$ and $j$ are computed as:

$$H'_{ij} = \frac{H_{ij}}{\sqrt{\hat{\mu_i} \cdot \hat{\mu_j}}} \cdot \hat{C}, \tag{S12}$$

where $\hat{\mu_i}$ and $\hat{\mu_j}$ are the predicted values from the model, and $\hat{C}$ is a rescaling factor.

#### Supplementary Note 3: Per-bin assembly refinement

The per-bin assembly refinement step in the METAHICT reassembly module adapts the read-recruitment and reassembly framework from [9]. In addition to the matched shotgun reads, METAHICT supplies EM-selected short-insert, shotgun-like intra-contig Hi-C read pairs as an additional read source. For each bin, reads are mapped to the bin's contigs with BWA MEM (v0.7.18, default parameters) [10] under two stringency presets: a strict preset (allowing no more than two mismatches) and a permissive preset (allowing up to five mismatches). A read pair is retained for the bin if at least one mate maps. Recruited reads from each preset are written to separate FASTQ files and assembled with SPAdes (v4.0.0; `-careful`) [11]. Contigs shorter than 500 bp are removed by default. CheckM2 (v1.1.0, default parameters)

[12] is applied to the original bin and to the two assemblies (strict and permissive) to estimate completeness and contamination. For each bin, a quality score defined as Score = Completion $-$ $k$ $\times$ Contamination is computed, where $k$ is a user-adjustable contamination penalty weight, set to 5 by default. The bin version with the highest score is retained as the best representative MAG.

### Supplementary Figures

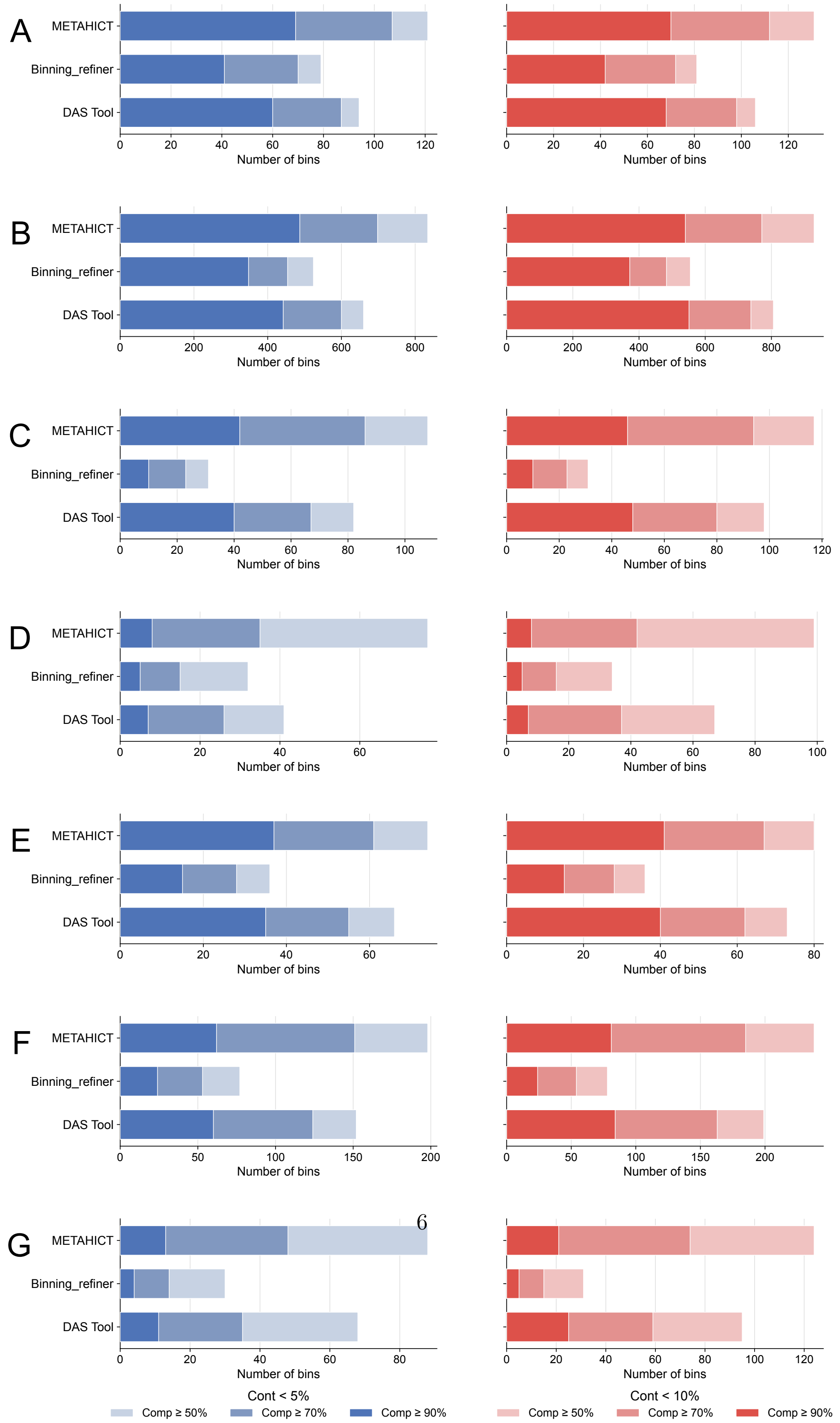

**Fig. S1: Comparison of ensemble binning strategies across benchmark datasets.** Numbers of MAGs meeting quality thresholds (contamination $< 5\%$ or $< 10\%$; completeness $\geq 50\%$, $\geq 70\%$, or $\geq 90\%$) for METAHICT, Binning_refiner, and DAS Tool in **A** human gut, **B** sheep gut, **C** pig gut, **D** cow rumen, **E** bovine skin, **F** wastewater, and **G** hydrothermal mats. The bin quality was assessed using CheckM2.

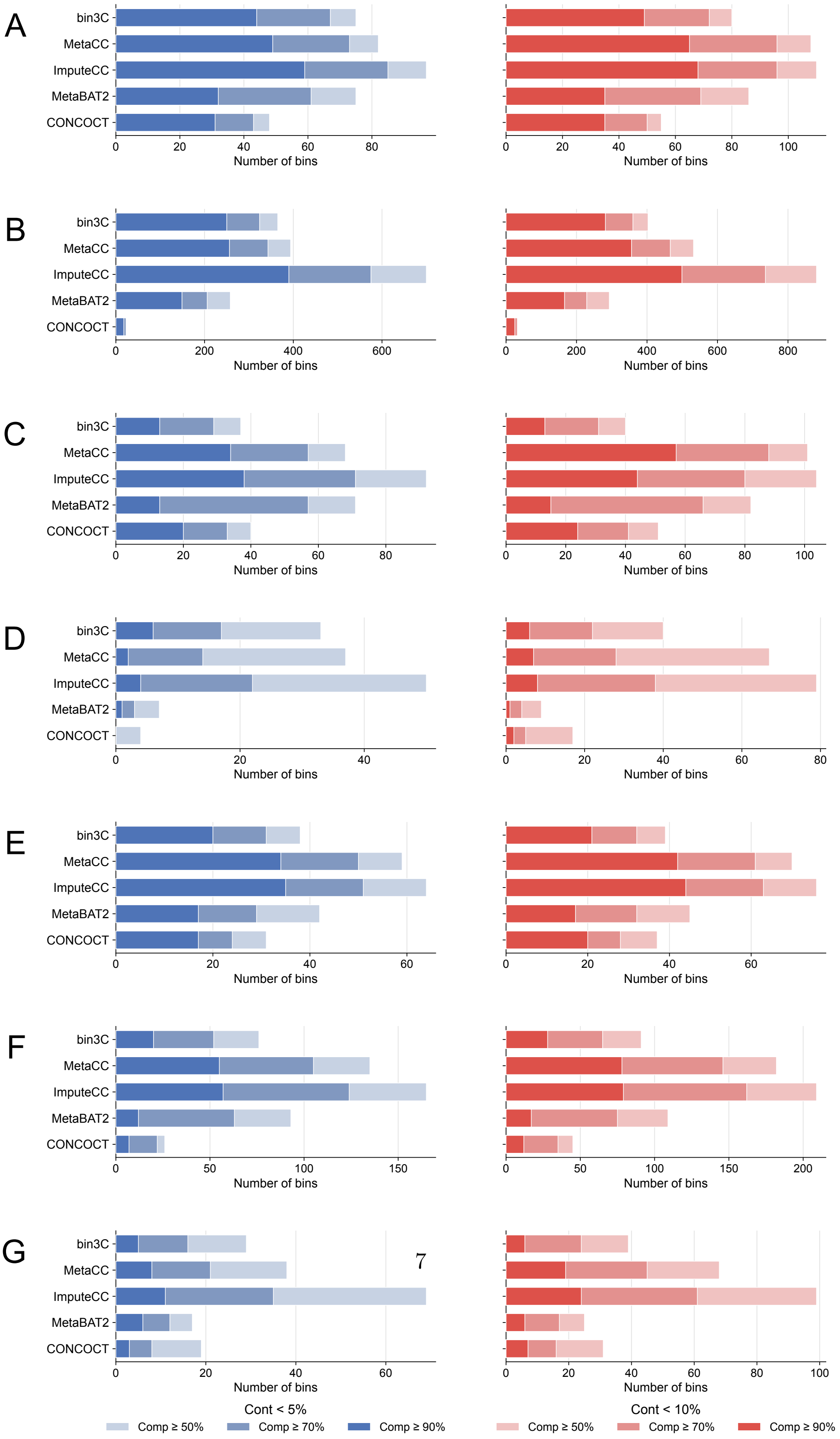

**Fig. S2**: **Benchmarking Hi-C-based and non-Hi-C binners across seven habitats.** Numbers of MAGs meeting quality thresholds (contamination $< 5\%$ or $< 10\%$; completeness $\geq 50\%$, $\geq 70\%$, or $\geq 90\%$) for bin3C, ImputeCC, MetaCC, MetaBAT2, and CONCOCT in **A** human gut, **B** sheep gut, **C** pig gut, **D** cow rumen, **E** bovine skin, **F** wastewater, and **G** hydrothermal mats. The bin quality was assessed using CheckM2.

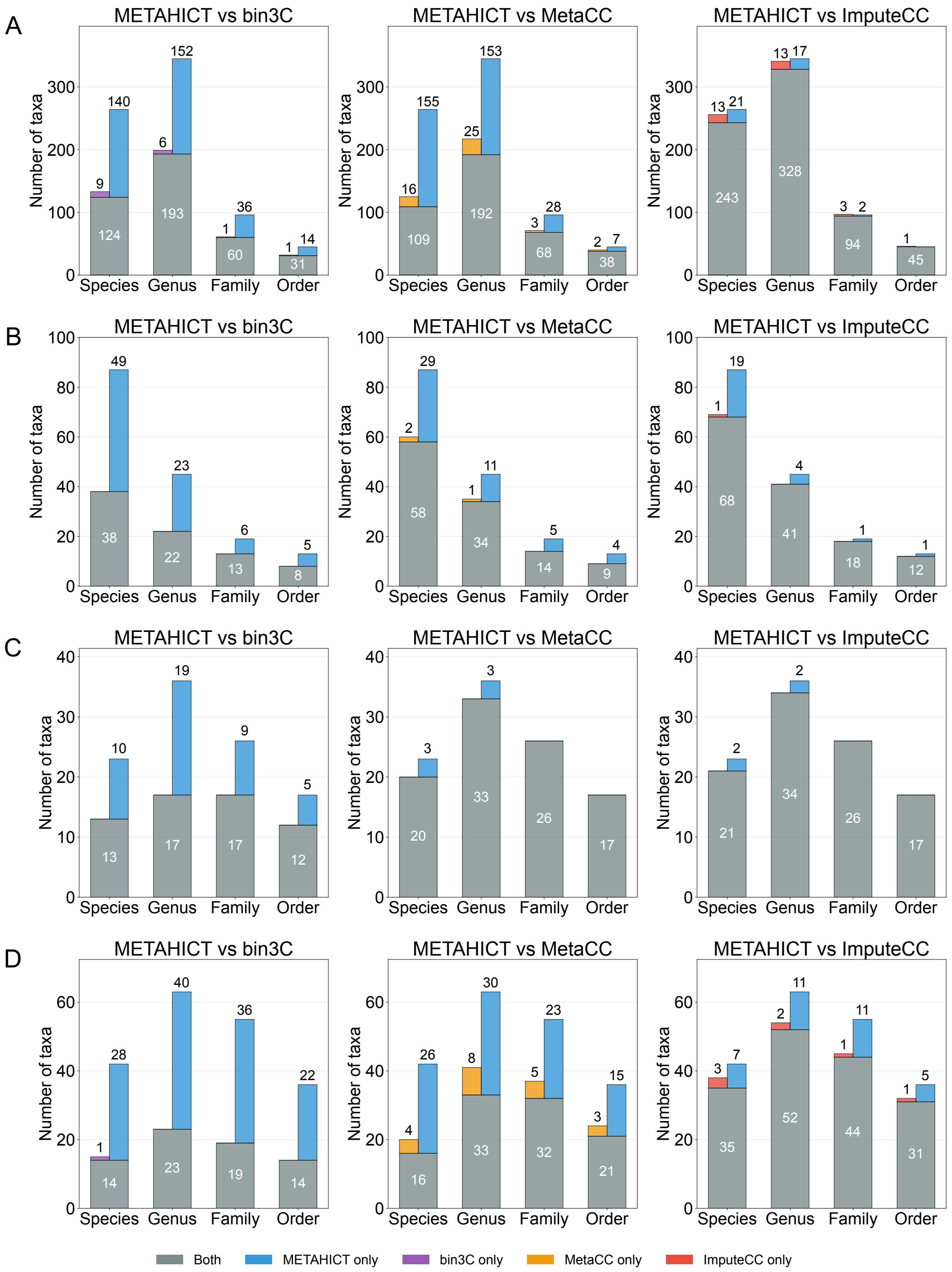

**Fig. S3**: **Taxonomic breadth of medium-quality MAGs.** Distinct GTDB-defined taxa (order, family, genus, species) recovered from medium-quality MAGs in pairwise comparisons between METAHICT binning and each Hi-C–based method (bin3C, MetaCC, ImputeCC). Bars are partitioned into taxa shared by both methods (grey) and taxa unique to one method (colors as indicated). Panels: **A** sheep gut, **B** cow rumen, **C** bovine skin, **D** hydrothermal mats.

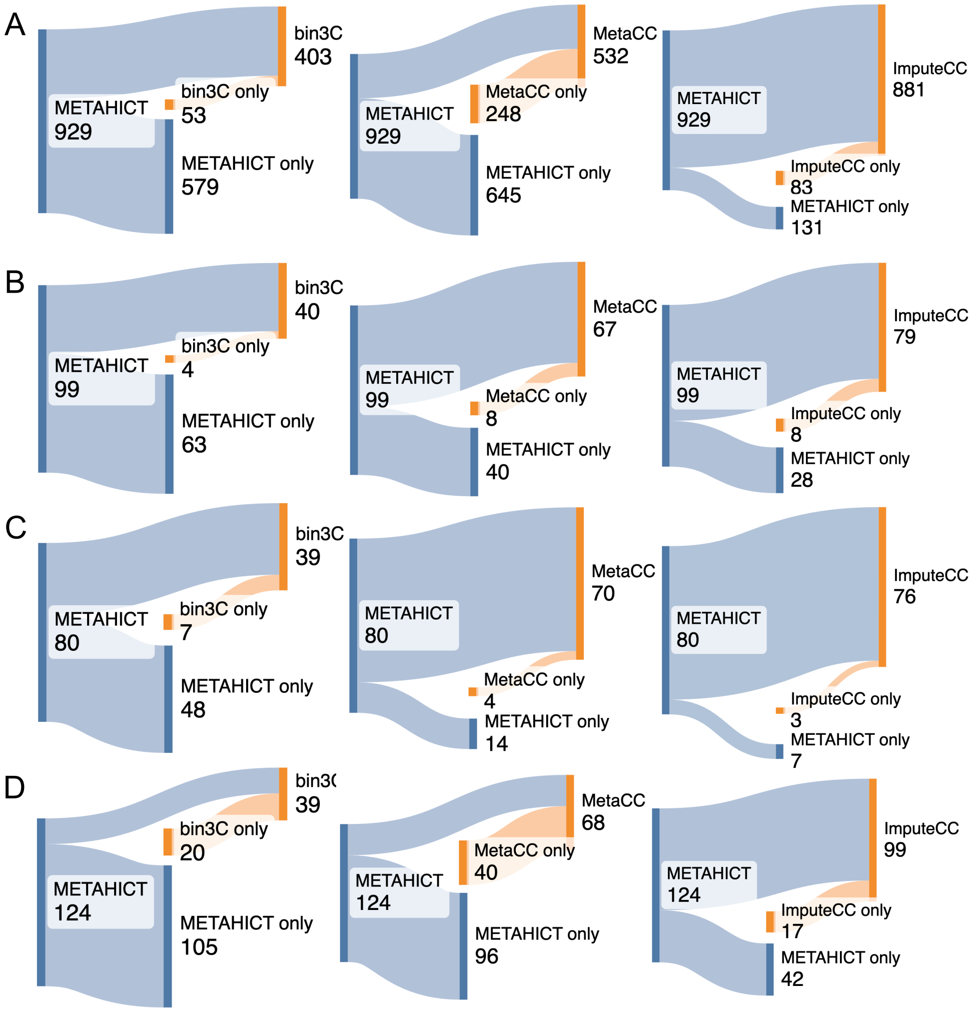

**Fig. S4**: **Overlap and uniqueness of medium-quality MAGs across metaHi-C datasets.** Sankey diagrams show the overlap of medium-quality MAGs (completeness $\geq$50%, contamination $<$ 10%) between METAHICT binning and each Hi-C–based method (bin3C, MetaCC, ImputeCC). Side bars represent all MAGs recovered by each method, while middle bars represent MAGs unique to each method. Panels: **A** sheep gut, **B** cow rumen, **C** bovine skin, **D** hydrothermal mats.

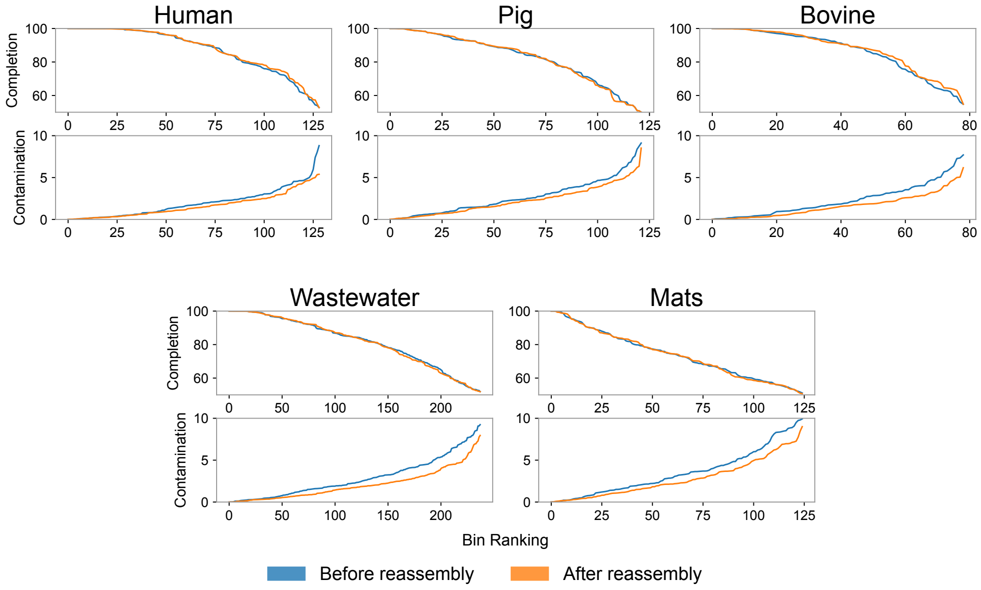

**Fig. S5**: **Bin quality curves before and after per-bin reassembly across five short-read datasets.** For each environment, bins are ranked by completeness (top panels) or contamination (bottom panels). Refinement lowers contamination curves, with the largest decreases observed for bins starting at higher contamination.

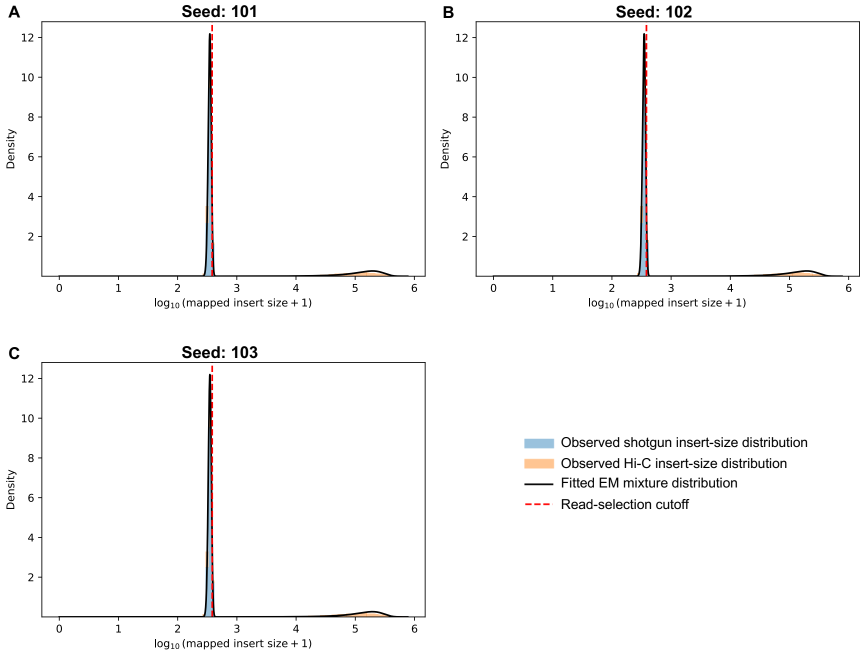

**Fig. S6**: **Controlled sim3C validation of top-contig EM fitting for short-insert read selection.** sim3C was used to generate metagenomic Hi-C reads from mock-community reference genomes with known abundances. Simulated reads were aligned to contigs assembled from the original mock-community shotgun library. The EM model was fitted using intra-contig read pairs mapped to the default 100 longest assembled contigs, and the resulting cutoff was applied to all intra-contig read pairs for evaluation. The histogram shows the top-contig mapped insert-size distribution used for EM fitting, with sim3C-derived shotgun and Hi-C labels shown separately. Shotgun-derived read pairs concentrate in the short-insert regime, whereas Hi-C-derived read pairs include a longer-insert tail.

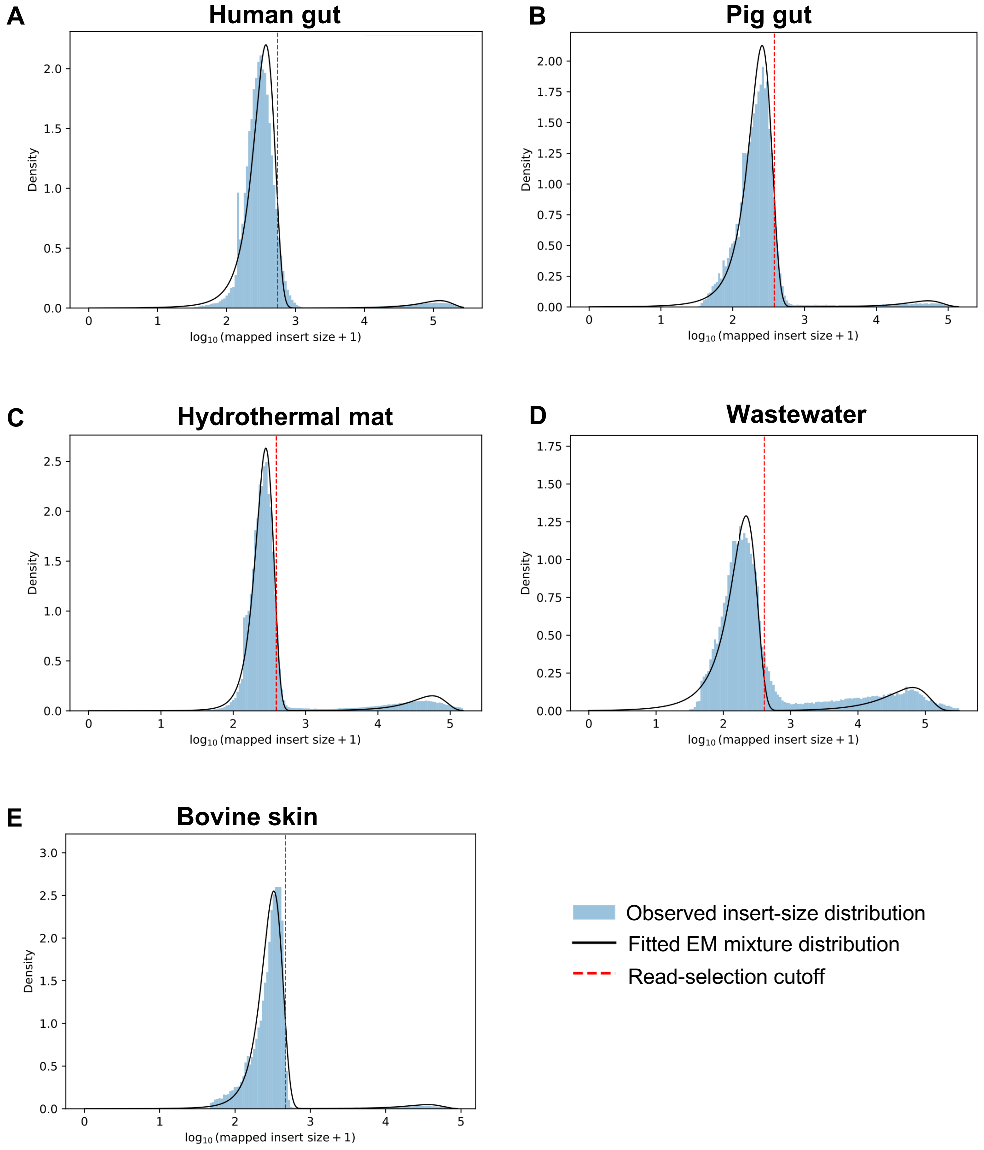

**Fig. S7**: **Real-dataset intra-contig mapped insert-size distributions and EM model fits.** For each short-read metaHi-C dataset, histograms show the mapped insert-size distribution of intra-contig read pairs associated with the 100 longest assembled contigs, black curves show the fitted EM mixture, and dashed vertical lines show the resulting read-selection cutoff. The fitted cutoff was applied to all filtered intra-contig read pairs for read selection. Visualization-only downsampling was used where necessary to reduce plotting density and was not used for EM fitting or selected-read fraction estimation. The empirical distributions contain dataset-specific shoulders and tails, supporting interpretation of the model as a pragmatic read-selection procedure rather than a complete generative model of molecular origin.

#### Supplementary Tables

**Table S1**: **Computational parameters used in this study.** Selective parameters central to the analyses described in the Methods are shown. The complete set of configurable parameters and default values is provided in the versioned configuration template and module documentation.

| Analysis step | Method or software | Parameter | Value used |
|---|---|---|---|
| Read preprocessing | BBDuk (BBTools v39.10) | Adapter and linker trimming | `ktrim=r`; `k=23`; `mink=11`; `hdist=1` |
| Read preprocessing | BBDuk (BBTools v39.10) | Quality and fixed-end trimming | `qtrim=r`; `trimq=10`; `ftm=5`; `ftl=10` |
| Read preprocessing | BBDuk (BBTools v39.10) | Minimum retained read length | `minlen=50` |
| Short-read assembly | MEGAHIT v1.2.9 | $k$-mer range and merge level | `k-min=21`; `k-max=141`; `k-step=12`; `merge-level=20,0.95` |
| Hi-C alignment | BWA-MEM v0.7.18 | Alignment options | `-5SP` |
| Hi-C alignment filtering | SAMtools v1.21 and METAHICT filtering | SAM flag, mapping-quality, and aligned-length filters | `-F 0x900`; MAPQ $\geq$ 30; aligned length $\geq$ 30 bp |
| Raw-contact construction | METAHICT | Minimum contact signal and contig length | 1 contact; 1,000 bp |
| Contact normalization | NormCC v1.2.0 | Spurious-contact filter | Lowest 5% of positive normalized contact values removed |
| Binning | CheckM2 v1.1.0 and adapted Binning_refiner | Minimum completeness and maximum contamination | 50% and 10%, respectively |
| Binning | Representative-bin score | Contamination penalty, $r$, in $S$ = completeness $- r \times$ contamination | 5 |
| EM model fitting | Two-component Gaussian mixture | Contigs used for fitting | 100 longest contigs |
| EM model fitting | Two-component Gaussian mixture | Component initialization | Lower 80% and upper 20% of mapped insert sizes |
| EM model fitting | Two-component Gaussian mixture | Convergence criterion | $\lvert\Delta \log L\rvert < 0.01$ or 100 iterations |
| EM read selection | Short-insert component | Cutoff quantile | 0.95 |
| Reassembly read recruitment | METAHICT | Strict and permissive mismatch cutoffs | 2 and 5, respectively |

Continued on next page

**Table S1 continued**

| Analysis step | Method or software | Parameter | Value used |
|---|---|---|---|
| Per-bin reassembly | SPAdes v4.0.0 | Assembly mode and minimum retained contig length | `careful`; 500 bp |
| Reassembly quality control | CheckM2 v1.1.0 | Contamination penalty | 5 |
| Contact-map visualization | METAHICT | Genomic bin size | 10 kb |
| Taxonomic annotation | GTDB-Tk v2.4.0 | Mode | `classify_wf` |
| MGE–host pairing | Within-sample standardization | $Z$-score threshold and minimum raw contact support | $Z > 0.5$; 2 contacts |
| Topology analysis | ccfind | Terminal fragment size, minimum identity, and minimum aligned length | 500 bp; 94%; 50 bp |

| Environment | Shotgun assembly type | Restriction enzymes | Hi-C library size |
|---|---|---|---|
| Human gut | Short | Sau3AI and MluCI | 25.9 Gbp |
| Sheep gut | Long | Sau3AI and MluCI | 32.3 Gbp |
| Pig gut | Short | HpyCH4IV | 29.8 Gbp |
| Cow rumen | Long | Sau3AI and MluCI | 10.1 Gbp |
| Bovine skin | Short | Sau3AI and MluCI | 39.2 Gbp |
| Wastewater | Short | Sau3AI and MluCI | 28.8 Gbp |
| Hydrothermal mats | Short | Sau3AI and MluCI | 41.6 Gbp |

**Table S2**: **Descriptions of seven environments used in the experiments.** In the 'Shotgun assembly type' column, 'Long' denotes assemblies generated from long-read shotgun data, whereas 'short-read' denotes assemblies generated from short-read shotgun data; all Hi-C libraries analyzed here were generated using short-read sequencing. The 'Restriction enzymes' column specifies the restriction enzymes used for constructing each corresponding Hi-C library.

| **Dataset** | **Number of contigs** | **Average length (bp)** | **Total length (bp)** |
|---|---|---|---|
| Human gut | 100,214 | 5,058 | 506,843,628 |
| Sheep gut | 47,246 | 90,466 | 4,274,155,803 |
| Pig gut | 197,665 | 2,979 | 588,749,094 |
| Cow rumen | 77,670 | 13,859 | 1,076,426,244 |
| Bovine skin | 93,480 | 3,132 | 292,741,964 |
| Wastewater | 677,757 | 2,570 | 1,742,003,537 |
| Hydrothermal mats | 520,858 | 2,227 | 1,159,854,252 |

**Table S3**: **Assembly statistics of contigs from each environment.** The sheep-gut and cow-rumen assemblies, which were generated from long-read shotgun data, have higher average contig lengths than the five short-read assemblies.

| Dataset | 3D ratio | Informative fraction |
|---|---|---|
| Human gut | 0.26 | 0.30 |
| Sheep gut | 0.03 | - |
| Pig gut | 0.29 | 0.32 |
| Cow rumen | 0.14 | - |
| Bovine skin | 0.17 | 0.20 |
| Wastewater | 0.47 | 0.59 |
| Hydrothermal mats | 0.43 | 0.50 |

**Table S4**: **Hi-C library indicators.** Summary of two QC metrics per dataset. The 3D ratio is computed in the alignment module, and the informative fraction is derived from the reassembly step. A dash indicates not applicable.

| Dataset | Best individual binner | Gain in qualifying MAGs, n (%) |
|---|---|---|
| **A. Near-complete MAGs** | | |
| Human gut | ImputeCC | +10 (16.9) |
| Sheep gut | ImputeCC | +97 (24.9) |
| Pig gut | ImputeCC | +4 (10.5) |
| Cow rumen | bin3C | +2 (33.3) |
| Bovine skin | ImputeCC | +2 (5.7) |
| Wastewater | ImputeCC | +5 (8.8) |
| Hydrothermal mats | ImputeCC | +2 (18.2) |
| **B. Medium-quality MAGs** | | |
| Human gut | ImputeCC | +21 (19.1) |
| Sheep gut | ImputeCC | +48 (5.4) |
| Pig gut | ImputeCC | +13 (12.5) |
| Cow rumen | ImputeCC | +20 (25.3) |
| Bovine skin | ImputeCC | +4 (5.3) |
| Wastewater | ImputeCC | +29 (13.9) |
| Hydrothermal mats | ImputeCC | +25 (25.3) |

**Table S5**: **Contribution of METAHICT bin-set consolidation relative to the best individual input binner.** For each dataset and MAG-quality category, the best individual comparator was the method recovering the largest number of qualifying MAGs among bin3C, MetaCC, and ImputeCC. Gain was calculated as the METAHICT count minus the best individual count, and the value in parentheses is the gain relative to the best individual count. The best comparator was selected independently for each dataset and quality category. All counts were obtained before per-bin reassembly. Near-complete MAGs have completeness $> 90\%$ and contamination $< 5\%$; medium-quality MAGs have completeness $\geq 50\%$ and contamination $< 10\%$.

| Dataset | Binning/refinement workflow reported in the original study | Numerical quality criterion reported in the original study | Reported MAG count | Reported taxonomic breadth |
|---|---|---|---|---|
| Human gut [13] | ProxiMeta [13] Hi-C deconvolution | CheckM [14] completeness $> 90\%$ and contamination $< 10\%$ | 50 | – |
| | | CheckM completeness $> 50\%$ and contamination $< 10\%$ | 75 | – |
| Sheep gut [15] | bin3C [3]; DAS Tool [16] was used for SCG-based quality scoring | DAS Tool SCG completeness $> 90\%$ and SCG contamination $< 10\%$ | 428 | 197 genera and 15 phyla |
| Cow rumen [17] | ProxiMeta and MetaBAT2 [18]; bin assignments were consolidated with DAS Tool | DAS Tool SCG completeness $\geq 90\%$ and SCG redundancy $< 5\%$ | 10 | – |
| | | DAS Tool SCG completeness $\geq 50\%$ and SCG redundancy $< 10\%$ | 103 | – |

**Table S6**: **MAG counts reported in the original publications of the benchmark datasets.** Only datasets for which explicit sample-specific MAG counts and corresponding numerical quality criteria were reported are shown. The numerical criteria, MAG counts, and binning or refinement workflows are reproduced from the original publications and were not recalculated using the METAHICT workflow. For the sheep-gut dataset, bin3C generated the bins and DAS Tool was used to calculate SCG-based quality metrics from the single bin3C bin set. For the cow-rumen dataset, DAS Tool consolidated the bin assignments generated by ProxiMeta and MetaBAT2. Because the original studies used different assemblies, binning workflows, quality-assessment methods, and reporting conventions, these values are provided as literature context rather than as a harmonized head-to-head benchmark. Counts reported under broader and stricter thresholds may overlap and should not be summed. Taxonomic breadth is included only when it was explicitly reported for the corresponding MAG collection; a dash indicates that it was not reported.

| Dataset | Contamination | | | Completeness | | |
|---|---|---|---|---|---|---|
| | Mean Δ | Raw $P$ | BH $q$ | Mean Δ | Raw $P$ | BH $q$ |
| **A. Overall reassembly effect: pre-reassembly versus METAHICT reassembly** | | | | | | |
| Human gut | -0.606 | $2.99 \times 10^{-11}$ | $\mathbf{9.97 \times 10^{-11}}$ | 0.546 | $6.62 \times 10^{-4}$ | **0.0011** |
| Pig gut | -0.626 | $7.57 \times 10^{-12}$ | $\mathbf{3.78 \times 10^{-11}}$ | -0.120 | 0.9214 | 0.9214 |
| Bovine skin | -0.787 | $1.64 \times 10^{-9}$ | $\mathbf{3.28 \times 10^{-9}}$ | 0.919 | 0.0031 | **0.0044** |
| Wastewater | -1.021 | $5.54 \times 10^{-24}$ | $\mathbf{5.54 \times 10^{-23}}$ | -0.210 | 0.6634 | 0.8149 |
| Hydrothermal mats | -0.761 | $1.25 \times 10^{-9}$ | $\mathbf{3.13 \times 10^{-9}}$ | -0.062 | 0.7334 | 0.8149 |
| **B. Incremental EM-selected-read effect: SG-only versus METAHICT reassembly** | | | | | | |
| Human gut | -0.215 | $6.94 \times 10^{-7}$ | $\mathbf{6.94 \times 10^{-6}}$ | 0.190 | 0.0072 | **0.0145** |
| Pig gut | -0.127 | 0.0017 | **0.0042** | -0.118 | 0.6291 | 0.6291 |
| Bovine skin | -0.213 | 0.0157 | **0.0261** | 0.680 | 0.0011 | **0.0037** |
| Wastewater | -0.139 | $1.24 \times 10^{-5}$ | $\mathbf{6.19 \times 10^{-5}}$ | 0.020 | 0.0744 | 0.1063 |
| Hydrothermal mats | -0.028 | 0.2750 | 0.3438 | -0.138 | 0.5038 | 0.5597 |
| **C. Shotgun-only effect: pre-reassembly versus SG-only** | | | | | | |
| Human gut | -0.391 | $1.86 \times 10^{-9}$ | $\mathbf{6.21 \times 10^{-9}}$ | 0.356 | 0.0469 | 0.0781 |
| Pig gut | -0.499 | $9.71 \times 10^{-12}$ | $\mathbf{4.85 \times 10^{-11}}$ | -0.001 | 0.9178 | 0.9178 |
| Bovine skin | -0.574 | $2.71 \times 10^{-5}$ | $\mathbf{5.43 \times 10^{-5}}$ | 0.239 | 0.3210 | 0.4013 |
| Wastewater | -0.882 | $1.27 \times 10^{-22}$ | $\mathbf{1.27 \times 10^{-21}}$ | -0.230 | 0.3033 | 0.4013 |
| Hydrothermal mats | -0.733 | $5.91 \times 10^{-8}$ | $\mathbf{1.48 \times 10^{-7}}$ | 0.076 | 0.3681 | 0.4090 |

**Table S7**: **Paired analysis of METAHICT reassembly and the EM-selected-read ablation.** Panel A compares pre-reassembly MAGs with the standard METAHICT reassembly procedure. Panel B compares SG-only reassembly with METAHICT reassembly and assesses the incremental contribution of the EM-selected short-insert Hi-C reads. Panel C compares pre-reassembly MAGs with SG-only reassembly. Mean changes (Δ) are calculated as the value after the indicated procedure minus the value before it; therefore, negative contamination changes and positive completeness changes indicate improvement. Each paired contrast included 131 matched MAG pairs from human gut, 117 from pig gut, 80 from bovine skin, 238 from wastewater, and 124 from hydrothermal mats. Statistical comparisons used paired two-sided Wilcoxon signed-rank tests. For each paired contrast, the raw $p$ values for contamination and completeness across the five datasets were adjusted together using the Benjamini–Hochberg procedure. Raw $p$ values and adjusted $q$ values are both reported; bold $q$ values indicate $q \leq 0.05$.

| Dataset | N50 before | L50 before | N50 after | L50 after |
|---|---|---|---|---|
| Human gut | 32,556 | 2,334 | 38,698 | 1,970 |
| Pig gut | 12,783 | 4,413 | 13,178 | 4,313 |
| Bovine skin | 14,171 | 2,304 | 18,848 | 1,654 |
| Wastewater | 8,061 | 18,356 | 9,878 | 14,321 |
| Hydrothermal mats | 4,983 | 14,497 | 5,681 | 11,755 |

**Table S8**: **Descriptive N50 and L50 statistics before and after per-bin reassembly.** Values summarize the contig-length distributions of the retained MAG collections before and after reassembly. Both summaries were calculated after retaining contigs of at least 1,000 bp. N50 and L50 are reported as descriptive statistics only.

| **Metric** | **Seed 101** | **Seed 102** | **Seed 103** |
|---|---|---|---|
| All intra-contig simulated read pairs | 4,597,877 | 4,601,453 | 4,600,854 |
| All shotgun-derived intra-contig read pairs | 3,567,688 | 3,569,994 | 3,572,271 |
| All Hi-C-derived intra-contig read pairs | 1,030,189 | 1,031,459 | 1,028,583 |
| Top-100-contig read pairs used for EM fitting | 1,113,714 | 1,113,878 | 1,111,807 |
| Top-100 shotgun-derived read pairs | 760,615 | 761,021 | 760,337 |
| Top-100 Hi-C-derived read pairs | 353,099 | 352,857 | 351,470 |
| Top-100 pairs / all intra-contig pairs (%) | 24.22 | 24.21 | 24.17 |

**Table S9**: **Read-pair statistics for the controlled sim3C validation.** sim3C-generated metagenomic Hi-C reads were simulated from mock-community reference genomes with known abundances using three random seeds. Simulated Hi-C reads were aligned to contigs assembled from the original mock-community shotgun library. The EM model was fitted using intra-contig read pairs mapped to the 100 longest assembled contigs, and the resulting cutoff was evaluated on all intra-contig read pairs. sim3C-provided shotgun/Hi-C labels were used for post hoc evaluation and were not used during EM fitting.

| Metric | Seed 101 | Seed 102 | Seed 103 |
|---|---|---|---|
| EM-selected read pairs | 4,019,919 | 4,022,824 | 4,023,167 |
| EM-rejected read pairs | 577,958 | 578,629 | 577,687 |
| EM-selected fraction of all intra-contig read pairs (%) | 87.43 | 87.43 | 87.44 |
| True shotgun-derived reads among EM-selected reads (%) | 84.15 | 84.14 | 84.17 |
| True Hi-C-derived reads among EM-selected reads (%) | 15.85 | 15.86 | 15.83 |
| Shotgun recall / shotgun retained fraction (%) | 94.81 | 94.81 | 94.80 |
| F1 score for shotgun-derived read pairs (%) | 89.16 | 89.16 | 89.17 |
| Overall accuracy (%) | 82.12 | 82.11 | 82.12 |

**Table S10**: **Controlled sim3C validation metrics for top-contig EM fitting.** The EM model was fitted using intra-contig read pairs mapped to the 100 longest assembled contigs, and the resulting cutoff was applied to all intra-contig read pairs for evaluation. Shotgun-derived intra-contig read pairs were treated as the positive class and Hi-C-derived intra-contig read pairs as the negative class. "True shotgun-derived reads among EM-selected reads" is equivalent to shotgun precision. "Shotgun recall" is equivalent to the fraction of all shotgun-derived intra-contig read pairs retained by the EM cutoff.

| Metric | Human gut | Pig gut | Bovine skin | Hydrothermal mats | Wastewater |
|---|---|---|---|---|---|
| All aligned Hi-C read pairs | 57,604,085 | 17,306,099 | 22,995,286 | 40,013,163 | 44,685,159 |
| Top-100-contig pairs used for EM fitting | 3,253,908 | 710,008 | 1,752,186 | 1,437,420 | 131,674 |
| EM-selected read pairs after applying cutoff to all intra-contig pairs | 39,315,915 | 11,204,243 | 18,510,629 | 21,144,519 | 21,427,067 |
| Selected / all aligned Hi-C read pairs (%) | 68.25 | 64.74 | 80.50 | 52.84 | 47.95 |

**Table S11**: **Real-dataset EM-selected short-insert read counts and fractions using top-contig EM fitting.** For each short-read metaHi-C dataset, the EM model was fitted using intra-contig read pairs mapped to the 100 longest assembled contigs. The resulting cutoff was applied to all intra-contig Hi-C read pairs. The table reports the number of aligned Hi-C read pairs, top-100-contig pairs used for EM fitting, EM-selected read pairs, and selected-read fractions relative to all aligned Hi-C read pairs.

| MAG | N50 before | L50 before | N50 after | L50 after |
|---|---|---|---|---|
| Bin32 | 15,016 | 26 | 17,774 | 24 |
| Bin39 | 20,692 | 32 | 25,294 | 27 |
| Bin66 | 50,324 | 12 | 63,128 | 9 |
| Bin88 | 44,425 | 10 | 49,752 | 8 |
| Bin25 | 45,676 | 9 | 52,082 | 8 |
| Bin70 | 46,925 | 16 | 62,001 | 12 |
| Bin15 | 54,110 | 10 | 63,725 | 8 |

**Table S12**: **Scaffolding contiguity statistics for seven *Faecalibacterium* MAGs from the human gut dataset.** N50 and L50 values are shown before and after YaHS-based Hi-C-guided scaffolding.

| Dataset | Total contigs | MGE contigs | MGE contigs with geNomad DTR/ITR | MGE contigs with ccfind terminal-overlap evidence | MGE contigs with both annotations | Non-MGE contigs with ccfind terminal-overlap evidence |
|---|---|---|---|---|---|---|
| Human gut | 169,633 | 4,544 | 50 | 45 | 42 | 96 |
| Sheep gut | 47,246 | 4,191 | 120 | 78 | 66 | 8 |
| Pig gut | 443,641 | 10,469 | 32 | 30 | 28 | 109 |
| Cow rumen | 77,670 | 10,468 | 7 | 14 | 3 | 16 |
| Bovine skin | 259,390 | 3,810 | 26 | 28 | 24 | 80 |
| Wastewater | 1,524,326 | 35,567 | 172 | 160 | 147 | 415 |
| Hydrothermal mats | 1,264,552 | 17,867 | 27 | 24 | 20 | 448 |

**Table S13**: **Contig-level sequence-topology evidence across the seven datasets.** The geNomad column reports MGE contigs annotated with direct terminal repeats (DTR) or inverted terminal repeats (ITR). The ccfind columns report contigs with terminal-overlap evidence consistent with a putatively circular assembly. The overlap column reports MGE contigs with both a geNomad DTR/ITR annotation and ccfind terminal-overlap evidence. These annotations provide sequence-topology evidence and are not interpreted as definitive confirmation of molecular circularity or used to modify CheckM2 completeness and contamination estimates.

| A. Read processing and contact construction | | | | | | |
|---|---|---|---|---|---|---|
| **Dataset** | **Metric** | **Preprocessing** | **Assembly** | **Alignment** | **Coverage** | **Contact** |
| Human gut | Time (h) | 0.69 | 1.79 | 0.50 | 0.77 | 0.12 |
| | Memory (GB) | 119.00 | 25.44 | 49.65 | 128.00 | 15.60 |
| Sheep gut | Time (h) | 0.48 | – | 1.57 | 1.92 | 0.12 |
| | Memory (GB) | 6.04 | – | 44.74 | 64.00 | 0.23 |
| Pig gut | Time (h) | 0.43 | 2.84 | 0.23 | 0.71 | 0.05 |
| | Memory (GB) | 5.90 | 18.51 | 19.52 | 128.00 | 5.68 |
| Cow rumen | Time (h) | 0.10 | – | 0.32 | 0.41 | 0.02 |
| | Memory (GB) | 5.71 | – | 40.83 | 62.20 | 0.24 |
| Bovine skin | Time (h) | 0.71 | 3.00 | 0.36 | 1.65 | 0.05 |
| | Memory (GB) | 6.04 | 39.33 | 26.60 | 128.00 | 7.18 |
| Wastewater | Time (h) | 1.58 | 13.60 | 0.82 | 3.73 | 0.13 |
| | Memory (GB) | 324.55 | 165.29 | 47.16 | 105.30 | 3.07 |
| Hydrothermal mats | Time (h) | 1.46 | 10.52 | 0.62 | 2.83 | 0.11 |
| | Memory (GB) | 292.25 | 161.13 | 44.15 | 128.00 | 2.00 |

| B. Genome reconstruction and downstream analysis | | | | | | |
|---|---|---|---|---|---|---|
| **Dataset** | **Metric** | **Binning** | **Reassembly** | **Scaffolding** | **Annotation** | **MGE** |
| Human gut | Time (h) | 1.20 | 6.43 | 0.84 | 0.61 | 1.16 |
| | Memory (GB) | 34.05 | 60.68 | 46.13 | 103.13 | 30.99 |
| Sheep gut | Time (h) | 29.82 | – | 0.63 | 0.93 | 2.07 |
| | Memory (GB) | 133.74 | – | 36.64 | 108.84 | 57.85 |
| Pig gut | Time (h) | 1.15 | 5.01 | 0.49 | 0.79 | 1.13 |
| | Memory (GB) | 114.12 | 30.31 | 118.05 | 96.67 | 5.00 |
| Cow rumen | Time (h) | 0.59 | – | 0.14 | 0.59 | 0.50 |
| | Memory (GB) | 28.80 | – | 20.80 | 102.25 | 16.72 |
| Bovine skin | Time (h) | 0.59 | 6.53 | 0.84 | 0.75 | 0.66 |
| | Memory (GB) | 13.40 | 72.51 | 128.00 | 96.53 | 4.50 |
| Wastewater | Time (h) | 3.43 | 11.97 | 0.52 | 0.88 | 3.72 |
| | Memory (GB) | 93.44 | 62.31 | 37.10 | 96.70 | 30.03 |
| Hydrothermal mats | Time (h) | 3.23 | 10.96 | 0.58 | 0.72 | 3.03 |
| | Memory (GB) | 130.30 | 67.00 | 37.00 | 94.49 | 19.54 |

**Table S14**: **Runtime and peak memory usage of METAHICT modules across benchmark datasets.** Runtime is reported in decimal hours and peak memory in gigabytes; both were obtained from module-level resource logs and rounded to two decimal places. Assembly and reassembly were performed only for the five short-read datasets. Measurements were obtained on an AMD EPYC 9654 96-core processor with 1,536 GB of memory, using up to 80 threads per workflow run. A dash indicates that the corresponding module was not run. The reported values are hardware- and configuration-dependent empirical resource estimates.
